# High-Frequency Focused Ultrasound Directly Activates the Auditory Midbrain and Protects Against Noise-Induced Hearing Loss in Guinea Pigs

**DOI:** 10.64898/2026.09.09.750439

**Authors:** Vinay Parameshwarappa, Guillaume Rastoldo, Emilie Franceschini, Olivier Macherey, Arnaud Norena

## Abstract

Although focused ultrasound (FUS) offers promising opportunities for neuromodulation, its ability to activate central auditory neurons remains uncertain. We targeted the central nucleus of the inferior colliculus (CNIC) in anesthetized guinea pigs using 20-MHz FUS while recording cochlear and cortical activity. Consistent with previous studies, FUS delivered through a small craniotomy evoked cortical responses through indirect cochlear activation, potentially mediated by skull mechanical coupling. However, under conditions that minimized cochlear interaction, FUS induced c-Fos expression in the CNIC and auditory cortex, suggesting direct central activation of the auditory afferent pathway. FUS additionally engaged the auditory efferent system, presumably through medial olivocochlear neurons. FUS delivered before acoustic trauma nearly prevented noise-induced hearing loss, limiting threshold shifts to <10 dB compared with ∼60 dB in controls. Gentamicin abolished both efferent modulation and protection, implicating cochlear efferent mechanisms. Together, these findings support FUS for auditory neuromodulation and cochlear protection against acoustic injury.

## Introduction

Neuromodulation, the ability to alter nervous system activity, offers promising opportunities for investigating neural circuits and treating neurological disorders^1^. Of the available approaches, focused ultrasound (FUS) is of particular interest as it can target deep brain structures non-invasively with high spatial precision. However, despite the growing interest in FUS neuromodulation, the underlying mechanisms, particularly the ability to directly activate central auditory neurons and modulate auditory pathways, remain poorly understood^2–4^. FUS is of particular interest for applications in the auditory system because of its potential to modulate neural activity at multiple levels of the auditory pathway. First, FUS neuromodulation could help interfere with and reduce the neuronal hyperactivity potentially associated with tinnitus^5–8^, in the same way that it could be used to reduce chronic pain^9^. Secondly, FUS could be used to restore hearing in people with severe hearing loss that cannot be improved with hearing aids. FUS could potentially provide targeted stimulation of the afferent (ascending) auditory pathway, either by targeting the cochlear spiral ganglion neurons using an implantable FUS-based cochlear device, or the inferior colliculus using non-invasive transcranial FUS. With sufficient high spatial and temporal resolution, FUS may be able to respect the spatial (tonotopic) and temporal coding of hearing information in a manner similar to cochlear or brainstem implants^10–12^. Thirdly, FUS could potentially be used to non-invasively activate the efferent (descending) auditory system which plays a role in hearing in noise and is involved in protection from acoustic overexposure^13–15^. However, despite growing interest in FUS neuromodulation, studies targeting the auditory system remain scarce. Furthermore, while it has been demonstrated that FUS (500 kHz) is capable of activating neurons in the somatosensory cortex of anesthetized mice in vivo^16^, such direct evidence is still lacking in the auditory system. A landmark article has shown that, when aimed at the exposed primary auditory cortex of guinea pigs, ultrasonic stimulation at 500 kHz does not directly activate cortical neurons, but is capable of activating the cochlea and then the afferent pathways all the way up to the auditory cortex^17^. A companion paper published the same year showed that this cochlear excitation induced widespread activation of the auditory cortex when transcranial ultrasound was focused on the visual cortex^18^. Notably, these studies employed low-frequency transducers that generate large focal volumes, resulting in broad ultrasound propagation through the skull and brain tissue in rodents^19,20^. Such widespread stimulation may facilitate intracranial standing waves^20^, leading to skull vibrations, bone conduction and/or cerebrospinal fluid (CSF) vibration capable of indirectly activating the peripheral auditory system^17,21^. Importantly, off-target auditory effects may also occur with transcranial FUS in humans, even when the ultrasound beam is spatially focused. Localized interaction between the ultrasonic field and the skull can generate shear waves within the skull that propagate through the bone, potentially reaching the cochlea and eliciting auditory responses via hair-cells activation^22^. Another more recent work showed a potential interesting long-lasting effect of 230 kHz ultrasound stimulation on the ABR amplitude. Indeed, ultrasound applied for 52 seconds was associated with a significant reduction in ABR amplitude for several hours or weeks^23^. However, the article does not specify whether or not the ultrasound may have stimulated the cochlea. This limitation is also present in a study that appears to show that prolonged stimulation (1 min) with ultrasound (1 MHz) increases the latencies of ABR waves^24^.

The conclusion to be drawn here is that, at this time, there is no direct, indisputable evidence of neural activation in the auditory system induced by ultrasound. Here, we addressed this question by using high-frequency FUS (20 MHz) to stimulate neurons in the central nucleus of the inferior colliculus (CNIC) via a small craniotomy in anesthetized guinea pigs. High-frequency FUS was chosen to enable spatially localized stimulation with submillimeter spatial resolution. This choice was further motivated by our recent demonstration that a 20-MHz FUS burst of 1-ms duration at acoustic pressures of 4-5.4 MPa can elicited calcium responses in neurons of the dorsal root ganglion^25^ and the spiral ganglion^26^. Given the pivotal position of the CNIC within the auditory pathway^27^, we asked whether and how FUS stimulation of the CNIC could modulate the ascending and/or descending auditory pathways. The CNIC is a major integrative hub of the auditory system where nearly all ascending auditory projections converge before relaying processed information to the thalamus and auditory cortex^28–30^. Second, since the CNIC is not located too high in the central auditory system, the CNIC "neural code" may be relatively simple and closely related to the spectro-temporal properties of sounds. In addition, the CNIC is a relatively large auditory structure with a well-defined tonotopic organization, potentially offering the possibility to restore hearing sensations as with cochlear or brainstem implants^27^. The CNIC is also the source of many efferent projections to the superior olivary complex and cochlea^31,32^, providing a pathway through which central auditory activity could modulate cochlear function.

Here, we show, consistent with previous studies, that FUS stimulation of the CNIC activates the cochlea ipsilateral to the stimulation and, to a lesser extent, the contralateral cochlea, likely through skull-mediated mechanical coupling. Importantly, c-Fos immunostaining revealed robust, spatially localized activation in the ipsilateral (i.e., FUS-stimulated) CNIC but not in the contralateral IC. The fact that this activation was also observed in a preparation minimizing cochlear activation demonstrates that FUS can directly activate the IC. c-Fos expression was also larger in the ipsilateral than in the contralateral auditory cortex suggesting that FUS engages the auditory afferent pathway. We further show that FUS stimulation of the CNIC also engages the auditory *efferent* system, as evidenced by reduced compound action potential (CAP) amplitudes and enhanced cochlear microphonic (CM) amplitudes in response to acoustic stimuli presented after FUS stimuli, both effects being an electrophysiological signature consistent with medial olivocochlear (MOC)-mediated modulation of cochlear function. These effects were abolished following pharmacological blockade of the MOC system with gentamicin. Finally, we examined the effects of FUS on noise-induced hearing loss (NIHL). This experiment was initially designed to investigate the extent to which noise-induced hearing loss might interfere with ultrasound-mediated cochlear activation but revealed an unexpected effect. Remarkably, FUS stimulation delivered before noise exposure almost completely prevented the 60-dB NIHL observed in control animals. This protective effect against noise trauma was only present when both cochlear and efferent pathways were concurrently activated and was abolished by gentamicin. This study provides the first evidence that FUS stimulation of the CNIC engages both the auditory afferent and efferent pathways. Furthermore, we demonstrate that activating the efferent system through FUS, together with cochlear excitation, can protect the cochlea from acoustic trauma to an unprecedented extent.

## Methods

### Animal preparation

All procedures involving animal care and use were approved by the Animal Care Committee of Bouches-du-Rhône, France and carried out in accordance with relevant guidelines and regulations. A total of 51 normal-hearing guinea pigs (500–1100 g) were used in this study (Supplementary Table 1).

All experiments were conducted under general anesthesia induced by intraperitoneal injection of ketamine (60 mg/kg) and medetomidine (0.5 mg/kg). To maintain anesthesia throughout the experiment, half doses of both agents were administered hourly.

Surgical procedures were performed as previously described^33,34^. Briefly, following anesthesia induction, a midline incision was made along the skull. The tissue overlying the frontal lobe was carefully removed using a scalpel blade to eliminate residual connective tissue and fascia. Four small screws were attached to the skull, and a larger central screw was fixed between them using dental cement to provide a mechanically stable anchor point for a metal head post. Subsequently, a small craniotomy was performed by removing the tissue, skull (≈0.25 cm^2^), and dura mater overlying the primary auditory cortex^35^ and visual cortex (10 mm caudal to bregma and 2.5 mm lateral to lambda). In order to better understand the mechanisms underlying FUS-evoked responses, the animal preparation was in some cases modified by using a larger craniotomy (∼1 cm²) or by sending FUS to the intact skull (c.f. Section **FUS stimulation through different craniotomy conditions and the intact skull**). Throughout the experiment, body temperature was maintained at ∼37 °C using a thermostatically controlled heating blanket and rectal probe. At the end of the experiment, animals were euthanized with a lethal dose of pentobarbital sodium (600mg/kg).

### Ultrasound stimulation

FUS was delivered using a focused ultrasound transducer with a center frequency of 20 MHz, a focal length of 1 2. 7 mm and a diameter of 6. 35 mm (V31 7-SM, Olympus). The FUS stimulus was generated by a function generator (33600B, Agilent, France) and amplified by a power amplifier (VBA1 00-30, Vectawave, UK). To calibrate the transducer, acoustic beam profiles were acquired using a pressure measurement system consisting of a 40-µm–diameter needle hydrophone (Model NH0040, Precision Acoustics, UK), a preamplifier, and a DC coupler. The hydrophone was mounted on a three-axis motorized translation stage (M403. 4PD, Physik Instrumente, Germany). After locating the center of the ultrasound focus, two-dimensional raster scans were performed in the elevation (X–Y) and azimuthal (Y–Z) planes over areas of 0. 6 × 0. 6 mm² and 3. 8 × 0. 8 mm², respectively. We found that the −6 dB lateral and axial beam widths were 0.22 mm and 2. 78 mm, respectively (Figures 1B and 1C).

**Figure 1.**
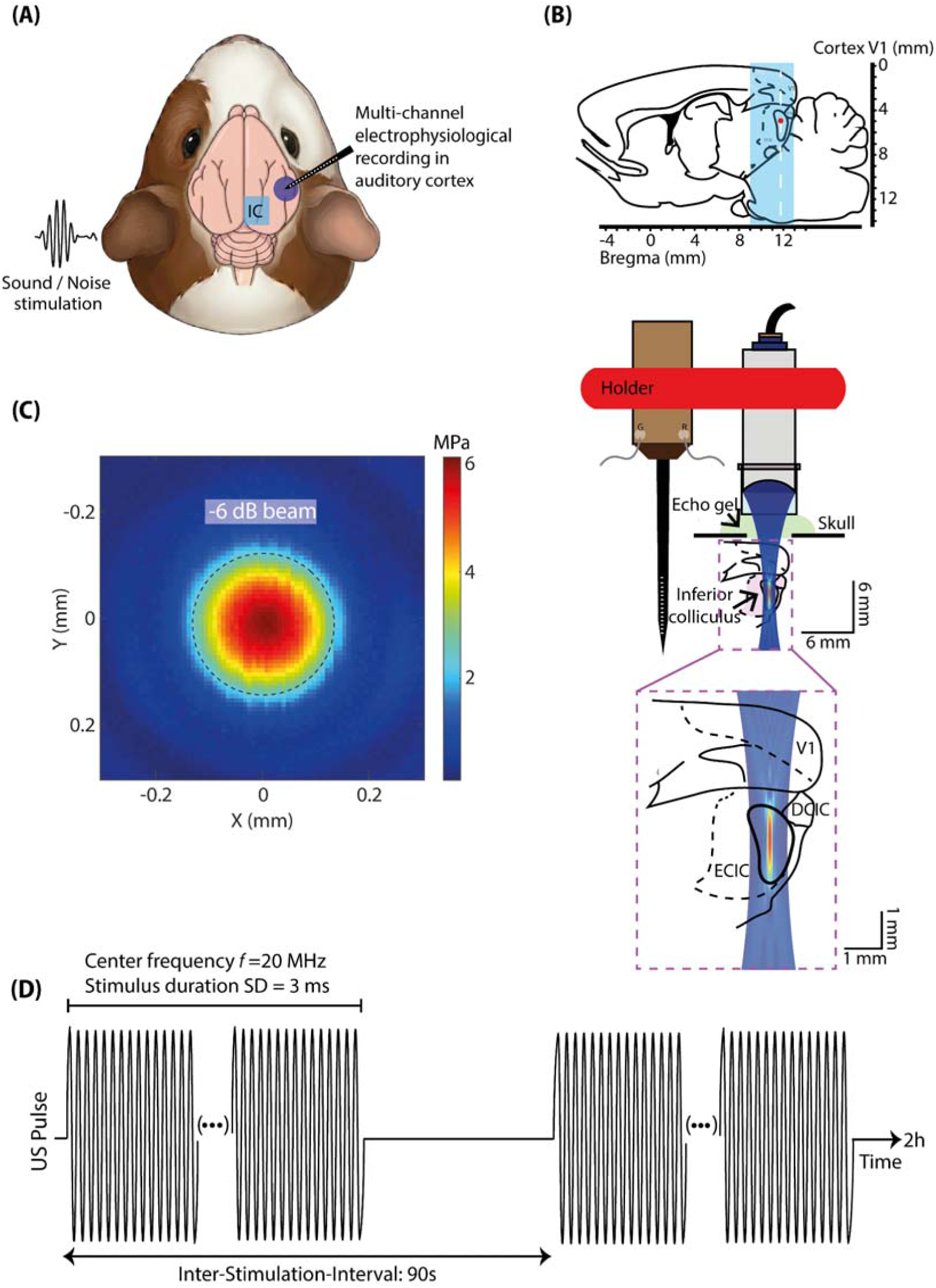
Experimental setup for high-frequency focused ultrasound (FUS) stimulation of the inferior colliculus (IC). (A) Schematic of the experimental configuration. High-frequency FUS was delivered to the right IC, acoustic stimuli were presented to the left ear, and neural activity was recorded from the right primary auditory cortex (A1) using a 16-channel linear electrode. (B) Schematic of the experimental FUS configuration showing transducer positioning, acoustic coupling with echo gel, and beam targeting of the Central Nucleus of the inferior colliculus (CNIC). The FUS transducer and 16-channel linear electrode were mounted in a custom 3D-printed holder with a 21 mm center-to-center separation. The enlarged view illustrates the ultrasonic beam trajectory and focal region relative to visual cortex (V1) and the dorsal and external cortices of the IC (DCIC and ECIC, respectively). (C) Two-dimensional map of the FUS pressure field at the focal plane, with the dashed contour indicating the −6-dB beam boundary. (D) Schematic of the FUS waveform and sonication parameters (20 MHz, 3 ms pulse duration, 6.8 MPa peak positive pressure, 90 s interstimulus interval) used for IC stimulation.

To achieve precise FUS targeting of the CNIC, we first located the CNIC electrophysiologically using a 16-channel linear microelectrode array^34^. A craniotomy was performed above the right CNIC, the microelectrode array was inserted into the CNIC and sound-evoked responses to tone pips were recorded to ensure the electrode array was in the CNIC (Supplementary Figures 1A and 1B). Following electrode removal, the FUS transducer was positioned such that its focal spot coincided with the previously recorded electrode location. This accurate positioning was achieved through a prior calibration procedure using a custom holder integrating the recording electrode and the FUS transducer (separated by 21 mm), which was translated along the X, Y and Z axes using a motorized micromanipulator (Shutter Instruments) (Figure 1B). A glass microbead (diameter of approximately 50-ym) was used as a common spatial reference. The electrode was advanced until it contacted the bead, and the corresponding 3D coordinates were recorded. The electrode was then withdrawn, and the transducer, operating in pulse-echo mode, was moved laterally until the echo from the bead was maximized, indicating the ultrasonic focus. Comparing the two sets of coordinates determined the displacement required to align the ultrasonic focus with the electrode position, thereby enabling accurate targeting of the CNIC with FUS. To minimize the amount of ultrasound gel required for acoustic coupling, the transducer was fitted with a 5-mm-long plastic coupling column filled with 1% agarose gel and coupled to the brain surface using a thin layer of ultrasound gel.

The FUS stimulus consisted of a series of 20-MHz sinusoidal burst signals with a peak positive pressure of 6.8 MPa and a peak negative pressure of -5.4 MPa. Each burst had a duration of 3 ms and consecutive bursts were separated by a gap of 90 s. The long interstimulus interval was chosen to minimize temperature rise in the CNIC and prevent transducer breakage. A total of 80 bursts were delivered during the two-hour FUS stimulation sequence. Spectral analysis of the acoustic waveform revealed harmonic components, with pressure amplitudes of 5.2 MPa at 20 MHz and 2.1 MPa at 40 MHz.

### FUS stimulation through different craniotomy conditions and the intact skull

Most experiments across the study were performed through a “small” craniotomy (“SC”) consisting of an opening of ∼ 0.5x0.5 cm = 0.25 cm^2^, with FUS delivered to the right IC (right-FUS), as described previously. This craniotomy was slightly smaller than the circular aperture of the 20-MHz transducer (∼6.35 mm in diameter), allowing the focused beam to pass through the craniotomy while retaining the possibility of ultrasound-skull interactions at the craniotomy edges, as the transducer could not be perfectly centered over the opening. To investigate the mechanisms underlying FUS-evoked responses, this preparation was modified in three different ways across the experiments. In some animals, FUS was delivered to the left (instead of the right) IC (n=3). In some others, FUS was delivered to the right IC through a large craniotomy (“LC”) of 1x1 cm=1 cm² (n=4). This LC condition prevented direct interactions between the skull and the FUS beam. Finally, in some other animals, FUS was applied directly onto the intact skull (“Skull”; i.e., without any craniotomy) (n=4), a condition expected to substantially limit 20-MHz US transmission into the brain due to strong attenuation and reflection at the skull. In this condition, the transducer was positioned so that its focus was located within the IC, as in the small- and large-craniotomy conditions.

### Auditory Brainstem Response (ABR) thresholds

Hearing thresholds of all animals were assessed using auditory brainstem responses (ABRs) as previously described^34^. Needle electrodes were placed subcutaneously at the skull vertex (active), behind the left or right mastoid (reference), and in the neck muscle (ground). Signals were amplified (10O×), bandpass filtered between 300 and 3000 Hz (Grass ICP 511 amplifiers), digitized, and averaged using a Micro1401 Plus system (Cambridge Electronic Devices, UK).

Auditory stimuli consisted of alternating-polarity tone pips (2-ms linear rise/fall, no plateau) presented monaurally at octave frequencies from 2 to 32 kHz via an in-ear miniature earphone at a repetition rate of 10 s⁻¹. Stimulus level ranged from 90 to 0 dB SPL in 10-dB steps. The number of stimulus repetitions depended on level, ranging from 100 repetitions at 90 dB SPL to 1000 repetitions at 0 dB SPL. ABRs were analyzed offline using custom MATLAB scripts. The five ABR waves correspond to sequential activation of brainstem nuclei, from the auditory nerve (wave I) to the region at or around the inferior colliculus (wave V).

Prior to averaging, individual ABR sweeps were screened for large-amplitude artifacts using an absolute voltage threshold of 25 µV. Sweeps exceeding ±25 µV at a given frequency and intensity were excluded together with their corresponding opposite-polarity sweeps. Artifact-free ABR sweeps were baseline-corrected by subtracting the mean voltage of a short reference segment, thereby centering the pre-stimulus baseline at 0 µV. For each subject and level, the sweeps were further averaged for analysis. The post-average residual noise was estimated for each animal and level using the across-sweep noise estimator method^36–38^. The across-sweep variance was calculated at seven time points ranging from 1 to 18 ms post-stimulus and averaged across these time points. The post-averaged residual noise variance was further computed as the averaged across-sweep variance divided by the number of sweeps. The wave amplitudes extracted from the ABR waveforms were assumed to be above noise floor whenever they exceeded +/- 2 standard deviations of the residual noise. Thresholds were defined as the lowest stimulus intensity at which a visually detectable ABR wave III response could be identified above the noise floor (wave III is the most stable and robust wave in the guinea pig)^39^. Thresholds (before and after noise exposure) were independently estimated for each animal by three observers (Authors VP, OM and AN), and the final threshold was defined as the average of these three estimates.

### Electrophysiology with microelectrodes

Electrophysiological recordings were performed in a sound-proofed booth using a linear 16-channel electrode array (100 µm spacing; A1 ×16-10mm-100-177-CM16LP, NeuroNexus, Ann Arbor, MI, USA) inserted successively in the right CNIC and in the right primary auditory cortex (A1), both localized using anatomical landmarks. Signals were acquired using a Tucker-Davis Technologies System 3 Pentusa (TDT, Alachua, FL, USA), amplified 10,000×, and bandpass filtered between 2 Hz and 5 kHz. Data were processed using a TDT System 3 Medusa multichannel acquisition system. Multi-unit activity (MUA) was sampled at 24,414.5 Hz and extracted from signals high-pass filtered at 300 Hz, whereas local field potentials (LFPs) were sampled at 1,061.5 Hz and extracted from signals low-pass filtered at 300 Hz, enabling simultaneous recording of spiking activity and LFPs.

First, neural recordings were made at the CNIC (except in animals of the “skull” group, i.e. with no craniotomy). The purpose of this step was to locate the CNIC so that the FUS transducer could be precisely positioned to ensure that the acoustic focus coincided with the CNIC (c.f. Methods Section: **Ultrasound stimulation**). The CNIC location and depth were verified by recording tone-pip–evoked responses (500 Hz–32 kHz, 1/8-octave steps, 70 dB SPL). The electrode was then removed, and the transducer was positioned at the same location over the x-axis with its acoustic focus targeting the CNIC (over the z-axis).

Subsequently, a linear 16-channel electrode was inserted perpendicularly into A1 using a Narishige microdrive. To localize tonotopic organization and cortical depth, responses were recorded to clicks, broadband noise bursts, and tone pips (500–32,000 Hz, 1/8-octave steps). Cortical depth was determined from the laminar profile of LFP responses to tone pips, characterized by short-latency, large-amplitude negative deflections in the middle cortical layers (approximately 500–800 µm).

Cortical responses were measured in response to both acoustic stimuli and FUS stimuli. Acoustic stimuli were generated in MATLAB and transferred to an RP2.1-based sound delivery system (Tucker Davis Technologies)^34^. Acoustic stimuli were presented from a calibrated free-field speaker (MF1, Tucker-Davis Technologies) positioned 10 cm from the ear. For sound-evoked measurements, the speaker was on the side of the ear contralateral to the cortex where the recordings were carried out. MUA and LFP tuning curves were recorded using 49 tone pips shaped by a gamma envelope and covering 8 octaves (with 1/8 octave step), from 0.5 kHz to 32 kHz, repeated 10 times. Tone pips were presented at different levels, from 70 to 20 dB SPL, with 10 dB step.

All electrophysiological data were first processed and analyzed using custom-written MATLAB routines. MUA or “spike events,” was detected by applying an amplitude threshold to high-pass filtered signals. The median of the negative values of the filtered signal was calculated, and the detection threshold was set to six times this median^40^. FUS-evoked cortical LFPs were quantified within a 100-ms window following each stimulus trigger. Peak latency was defined as the time of the maximal negative deflection occurring within 10–50 ms after trigger onset. Sound-evoked frequency tuning curves were derived from both MUA and LFPs. For MUA, 1-ms bin post-stimulus time histograms (PSTHs) were generated for each gamma tone pip, and spikes occurring within 0–100 ms after stimulus onset were averaged across repetitions at each intensity to generate frequency–intensity response profiles. LFP frequency–intensity response areas were constructed from trial-averaged traces by extracting the largest negative deflection within 50 ms after stimulus onset. Tuning curves from individual animals were aligned to their characteristic frequency (CF; the frequency with the lowest response threshold), expressed in octaves relative to CF, and then averaged across animals. Cortical threshold was defined as the stimulus intensity eliciting 25% of the maximal response. Current source density (CSD) analysis was performed as described previously^41^. Briefly, averaged translaminar LFPs were smoothed using a three-point Hamming window, and one-dimensional CSD was estimated from the second spatial derivative of the LFP across cortical depth^42^:

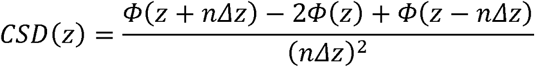

where Φ here is the LFP, *z* is the spatial coordinate, Δ*z* is the interelectrode spacing (Δ*z* = 100 μm) and *n* is the differentiation grid.

### Cochlear responses

The response of the cochlea to both FUS-mediated activation and acoustic stimuli was assessed using compound action potentials (CAPs), reflecting the summed activity of the cochlear nerve^43–45^. Recordings were obtained using a custom-made ball-contact Platinum electrode positioned at the cochlear round window (RW)^46^. Animals were anesthetized as described above and placed on a heating blanket in a sound-attenuated chamber. After shaving the surgical site, the bulla was approached from behind the pinna and drilled (∼1-mm² opening) to expose the RW membrane. The electrode was positioned in the RW niche, and correct placement was verified by recording CAP responses to brief acoustic clicks. The electrode was then sealed with dental cement to stabilize it throughout the experiment. Signals were amplified (20,000×; AC-coupled differential amplifier, 100 Hz–10 kHz; Grass ICP 511) and digitized via a sound card (RME Babyface Pro FS) and CED 1401 data-acquisition system.

To assess cochlear activation induced by FUS, CAP responses were recorded in either the right (ipsilateral) cochlea in 2 SC and 2 “Skull” animals or the left (contralateral) cochlea in 4 SC and 2 “Skull” animals during stimulation of the right CNIC. In addition (for comparison), acoustically evoked CAPs were recorded in response to alternating-polarity tone pips (4, 8, and 16 kHz; 40–80 dB SPL; 50 repetitions per condition).

### Effect of FUS on cochlear responses (CAP and Cochlear microphonics) to sound stimuli

To assess whether FUS induced sustained modulation of cochlear function, tone-evoked CAPs and Cochlear Microphonics (CMs), which predominantly reflects outer hair cell activity^41,43^ were recorded from the cochlea contralateral to the FUS-stimulated inferior colliculus at two frequencies (4 and 8 kHz; 2-ms linear rise/fall, no plateau; alternating polarity; 40–80 dB SPL). Stimuli were presented every 20 ms (50 Hz), and responses were averaged across 30 repetitions. This measure was performed in a total of 9 animals from different preparations, including SC (n=4), LC (n=3) as well as two SC animals treated with Gentamicin (c.f. Section **Pharmacological disruption of efferent cholinergic transmission using gentamicin).**

The CAPs and CMs were collected before FUS stimulation (−60 and −2 min), throughout the 2-hour FUS stimulation sequence (during the 90-s FUS inter-burst intervals), and up to 1 h after FUS stimulation. This means that sound and FUS stimuli were always presented non-simultaneously. Tone pips were delivered through an in-ear miniature earphone. CAPs were obtained by averaging responses to alternating-polarity stimuli, and amplitudes were quantified as the N1–P2 peak-to-trough difference. CMs were isolated by subtracting responses to opposite stimulus polarities, and amplitudes were quantified as the root mean square (RMS) voltage over the stimulus duration.

### Acoustic trauma

Guinea pigs were exposed to acoustic trauma after the FUS stimulation sequence. The effects of FUS on noise-induced hearing loss was studied in different groups: 1) FUS applied through a small craniotomy targeting the right CNIC (n = 6), 2) FUS applied through a small craniotomy targeting the left CNIC (n = 3), 3) FUS applied through a large craniotomy targeting the right CNIC (LC; n = 4), 4) FUS applied on the intact skull overlying the right CNIC (Skull; n = 4), 5) FUS delivered in animals treated with Gentamicin through a small craniotomy targeting the right CNIC (GM; n=2), and 6) control animals that received no FUS (n = 5). Control animals remained in silence for the corresponding 2-h pre-trauma period. Acoustic trauma consisted of a continuous 8-kHz pure tone at 115 dB SPL for 1 h, delivered in a sound-attenuated booth through a speaker positioned 15 cm from the left ear. ABRs and sound-evoked A1 tuning curves were recorded before and after noise exposure to assess functional protection. ABRs and sound-evoked A1 tuning curves were recorded in all groups before and after noise exposure to assess functional protection or recovery from acoustic trauma.

### Pharmacological disruption of efferent cholinergic transmission using gentamicin

To block cholinergic transmission at outer hair cells, gentamicin (GM; 250 mg/kg) was administered intraperitoneally in two SC animals^47,48^. CAP and CM responses were recorded before and 2 h after GM administration. FUS stimulation was initiated 2 h following GM injection, and changes in CAP and CM responses were monitored throughout and after the FUS stimulation period, as well as following noise trauma. Cortical tuning curves were also recorded before and after noise exposure to assess whether FUS-induced protection persisted after GM administration.

### Assessment of FUS-induced c-Fos expression across the auditory pathway

FUS-induced c-Fos expression was assessed in different experimental conditions involving a total of 6 animals: (1) no sound and no FUS stimulation (negative control; n = 1); (2) white-noise stimulation (100-ms duration, 60 dB SPL, presented every 5 s) delivered to the right ear and no FUS stimulation (positive control; n = 1); (3) FUS stimulation through LC condition (n = 2); and (4) FUS stimulation through SC condition (n = 2). The sample size used in the present study is based on earlier studies^49,50^. To minimize the influence of background acoustic stimulation on neural activity, the external ear canals were filled with dental cement before surgery. Following completion of all surgical procedures, including electrode implantation in the IC and A1 and placement of the ultrasound transducer over the IC, animals were maintained in silence for 3 h before FUS stimulation to minimize surgery- and sound-induced c-Fos expression unrelated to ultrasound stimulation^51^. Following the experimental protocol, animals remained undisturbed in silence for an additional 45 min to allow robust c-Fos expression before transcardial perfusion and tissue fixation.

Anaesthetized guinea pigs were transcardially perfused using 0.9% saline solution (NaCl), followed by 4% paraformaldehyde (PFA). Whole brains were dissected from the skull and postfixed overnight in fresh PFA, then rinsed three times for 5 min in PBS. Prior to sectioning, the brains were cryoprotected by successive transfers into increasing concentrations of sucrose solution (10%, 20%, and 30% in 0.1 M PBS) for 72 h at 4 °C, until the tissue had fully sunk. Brains were rapidly frozen with pentobarbital for 5 minutes and coronally sectioned at 40 μm using a Leica CM3050 freezing cryostat.

Floating brain sections of guinea pigs were washed (2 x 5 min) with PBS in multi-well plates. Blocking was done by incubation (90 min) in 10% Donkey serum and 0.5% Triton X-100. Slices were incubated 72h at 4°C with the following primary antibodies: rabbit anti-C-fos (1:1000, Sigma Aldrich, ABE457) and mouse anti-NeuN (1:500, Sigma Aldrich, MAB377). Sections were then rinsed in PBS and incubated for 2h at room temperature with fluorescent secondary antibodies used as follows: Alexa Fluor 594 donkey anti-mouse (1:500), Alexa Fluor 488 goat anti-rabbit (1:500). Sections were incubated with the nuclear marker DAPI (1:10 000, Invitrogen, D1306) for 3 minutes. Finally, brain sections were mounted onto SuperFrost/Plus glass slides and air-dried before being mounted with Roti®Mount FluorCare antifade reagent.

### Assessment of FUS-induced cell death in the inferior colliculus

Three hours after the end of FUS stimulation, guinea pigs (n=3 with FUS targeting the right CNIC, n=3 without FUS) were deeply anesthetized and transcardially perfused with 0.9% saline (NaCl) followed by 4% paraformaldehyde (PFA), as described above for c-Fos evaluation. Floating brain sections were processed using the same protocol as described above and incubated overnight at 4°C with primary antibody against cleaved caspase-3 (1:300; CST, #9664), followed by Alexa Fluor 488-conjugated goat anti-rabbit secondary antibody (1:500) and DAPI (1:10 000, Invitrogen, D1306). To evaluate cell death, fluorescence intensity was quantified using ImageJ from 10 single-plane images (425 µm²) per animal, sampled across rostrocaudal levels of the ipsilateral IC, in control (no FUS) and FUS-stimulated animals.

### Fluorescence image acquisition and quantification

c-Fos quantification was achieved for the six animals, including 2 control (1 negative, 1 positive), 2 SC, and 2 LC. Fluorescence images were acquired using a Zeiss LSM 710 NLO confocal microscope with a 20× Zeiss Plan Apochromat objective (NA 0.8, DIC M27), controlled with Zen software (ZEN 2012 Black edition, Carl Zeiss Microscopy GmbH, Germany). Z-stacks (6–13 optical sections, 0.95 µm step size) were acquired to capture c-Fos labeling across the auditory pathway (A1, IC, and DCN). Images were acquired at a resolution of 1024×1024 pixels at subsaturating laser intensities for each channel. C-Fos-positive cells were counted using ImageJ (Cell Counter plugin) within a region of interest of 425 µm², delineated using an integrated microscopic counting chamber, in 1 section in IC and 10 sections in DCN and A1. In IC, the c-Fos positive cells were counted only in the section where the c-Fos immunostaining was maximum. To illustrate the region targeted by FUS stimulation in the IC, multipanel tile images were acquired and stitched to encompass the full extent of c-Fos-activated cells; the FUS-targeted region was identified by the electrode scar within the IC.

For cell death quantification in the IC, single-plane images were acquired at a resolution of 1452×1452 pixels using a 63× Zeiss Plan Apochromat objective (NA 1.40, DIC M27). Mean fluorescence intensity of cleaved caspase-3 was quantified using ImageJ within a region of interest of 212 µm², delineated using an integrated microscopic counting chamber. Ten sections per animal were analyzed for control and FUS-stimulated animals (n = 3/group).

## Statistical Analysis

### Effect of FUS on tone-evoked CAP and CM

CAP and CM amplitudes were analyzed using generalized linear mixed-effects models (GLMMs), with Time (pre-versus post-FUS), Frequency (4 and 8 kHz), Intensity (40–70 dB SPL), and stimulation condition (small-craniotomy right-IC FUS, large-craniotomy right-IC FUS, and gentamicin-treated small-craniotomy right-IC FUS) as fixed effects and animal as a random intercept. Five candidate models of increasing complexity were compared, and the best-fitting model was selected based on the corrected Akaike information criterion (AICc). AICc weights were used to assess relative model support (Supplementary Table 2). Analyses were performed in R using the lme4 package, with AICc values calculated using MuMIn.

### Effect of acoustic trauma on ABR thresholds and Wave III amplitudes

Noise-induced threshold shifts were calculated as the difference between pre- and post-exposure ABR thresholds. A linear mixed-effects model was used to evaluate ABR threshold changes. Fixed effects included Time (before and after noise trauma), Frequency (2, 4, 8, 16, and 32 kHz), and Group (Control, SC: Ipsi US, LC: Ipsi US, SC: GM+Ipsi US, Skull: Ipsi US, and SC: Contra US), and animal was included as a random intercept.

ABR input-output growth functions were derived from ABR wave III amplitude measured at 4 and 8 kHz across stimulus levels, ranging from 80 to 40 dB SPL both before and after noise exposure. Wave III amplitude was quantified using a peak-to-trough method, defined as the voltage difference between the largest positive deflection occurring 2.5–4.5 ms after stimulus onset and the subsequent negative trough occurring within 4.0–5.5 ms after stimulus onset. As for the CAP and CM analysis, several GLMMs were fitted to these data. Fixed effects included Time (before vs after noise trauma), Frequency (4 and 8 kHz), Intensity (70–80 dB SPL), and Group (Control, SC: Ipsi US, LC: Ipsi US, SC: GM+Ipsi US, Skull: Ipsi US, and SC: Contra US), with animal included as a random intercept.

### Effect of acoustic trauma on Cortical Frequency tuning

Statistically significant changes in tuning curve profiles after noise exposure were assessed using non-parametric cluster analysis^52,53^ as described previously^41^. Briefly, for each frequency–intensity pair, t scores were calculated for the difference between post- and pre-noise exposure profiles, producing a 2-D t-map. Non-significant t values were set to zero, and contiguous t values of the same sign were clustered, with cluster statistics defined as the sum of absolute t scores. Significance was evaluated via a bootstrap procedure (cluster α < 0.05, two-tailed α = 0.05), generating 1,000 random partitions by shuffling pre- and post-exposure data with replacement. For each shuffled dataset, 2-D t maps and cluster statistics were computed to create a null distribution. Clusters in the actual data exceeding the 95th percentile of the bootstrap distribution were considered significant.

### Three-dimensional acoustic and thermal simulations

To estimate the acoustic pressure reaching the IC and assess the potential thermal effects associated with FUS stimulation, nonlinear ultrasound wave propagation was simulated using the k-space pseudospectral method-based solver, k-Wave (k-Wave Version v1.4.0), accounting for compressional wave propagation and acoustic absorption in heterogeneous media (water, brain and bone), but not mode conversion^54^. The experimental 20-MHz transducer (Olympus V317-SM; 6.32-mm aperture, 12.7-mm radius of curvature) was modeled as a single spherically focused bowl element. Its curved emitting surface was represented using the off-grid source formulation implemented in kWaveArray, which uses band-limited interpolation to minimize staircasing errors associated with curved sources on Cartesian grids^55,56^. The simulated signal consisted of a 20-cycle burst in homogeneous water or 50-cycle burst in heterogenous media. The computational grid spacing was 12.5 µm, corresponding to J/6 at 20 MHz and J/3 at 40 MHz, where J is the wavelength in water. Detailed acoustic and numerical parameters are provided in Supplementary Table 3.

The simulations were first performed in homogeneous water and compared with the amplitudes of the fundamental (A1) and second-harmonic (A2) components measured experimentally with the hydrophone. The input pressure of the simulated transducer was then adjusted to produce similar pressure amplitudes (A1 and A2) at the US beam focus in water as those obtained with the real transducer. The same input pressure was then applied on the surface of the transducer for the simulation of the US propagation through the guinea pig tissues. The three-dimensional guinea-pig skull geometry was obtained from a micro-computed tomography (µCT) dataset of *Cavia porcellus* (TMM M-7283; DigiMorph, University of Texas at Austin) with spatial resolution of 41 µm. The CT-derived skull data were registered to the coordinate system of the experimental craniotomy and the ultrasound beam, then resampled onto a 12.5-µm isotropic computational grid. Because modeling the entire guinea pig head at this resolution would have exceeded the available computational memory, the simulations were performed using a unilateral head model restricted to the ultrasound propagation pathway in a domain of 1701 × 960 × 1944 voxels. The computational domain encompassed the transducer aperture, craniotomy, inferior colliculus, and ipsilateral cochlear landmarks, excluding the contralateral side, which lay outside the experimental acoustic propagation pathway. Spatially heterogeneous maps of sound speed, density, acoustic attenuation, and the nonlinearity parameter were assigned to the different tissues^57,58^ (Supplementary Table 3). Peak-positive and peak-negative pressures were recorded throughout the computational domain.

To estimate FUS-induced heating, the three-dimensional acoustic pressure field and material segmentation were resampled to 25-µm isotropic resolution and used in a heterogeneous bioheat model. Temperature evolution was calculated using the Pennes bioheat equation^59^:

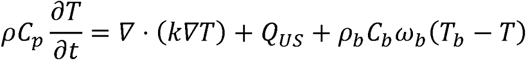

where *p*, *C_p_*, and k denote material-specific density, heat capacity, and thermal conductivity, respectively, and *Q_US_* represents acoustic heat deposition calculated as *Q_US_* — *ap*^2^/*Z*, where o is the absorption coefficient, *p* is the simulated acoustic-pressure amplitude, and *Z* is the acoustic impedance^60,61^. The last term corresponds to the perfusion process, where zu *ω_b_*, *p*, *C*,and *T_b_* are the blood perfusion rate, blood density, blood specific heat and blood ambient temperature, respectively. The model incorporated heterogeneous tissue properties, brain perfusion, zero-normal-heat-flux outer boundaries, a 3-ms FUS exposure, and 200-ms post-stimulation cooling. Detailed thermal parameters are provided in Supplementary Table 3.

Model construction and postprocessing were performed in MATLAB, and the large-scale US propagation simulations were executed on a Linux high-performance computing node using a parallel OpenMP implementation of the k-Wave solver with 64 threads (kspaceFirstOrder-OMP solver).

## Results

### Indirect activation of the cochlea and the ascending auditory pathway by high-frequency FUS stimulation

We first studied whether high-frequency FUS can directly activate afferent neurons in the CNIC and drive activity along the ascending auditory pathway in anesthetized guinea pigs. Local field potentials (LFP) and multi-unit activity (MUA) were recorded from the primary auditory cortex (A1) using a linear 16-channel microelectrode array while stimulating the CNIC with a 20-MHz FUS transducer (Figure 1A) (see methods). To achieve precise FUS targeting of the CNIC, we first localized the CNIC electrophysiologically using 16-channel linear microelectrode array. A small craniotomy (SC) was performed above the right CNIC, the microelectrode array was inserted into the CNIC and sound-evoked responses to tone pips were recorded to ensure the electrode array was in the CNIC. CNIC localization was confirmed by the presence of robust, short-latency and time-locked sound-evoked LFP responses (Supplementary Figures 1A & 1B), consistent with previous electrophysiological recordings from this structure. The electrode was then removed, and the FUS transducer was positioned using a pre-calibrated electrode-transducer holder, such that the acoustic focus coincided with the previously identified CNIC location, approximately 4–5 mm below the cortical surface (Figure 1B). The 20-MHz frequency was selected to achieve a tightly focused acoustic beam, with measured −6-dB lateral and axial beam widths of 0. 22 and 2. 78 mm, respectively (Figures 1B & 1C). FUS stimulation consisted of 3-ms pulses delivered at 90-s inter-stimulation intervals with a peak positive pressure of 6.8 MPa (Figure 1D). These FUS parameters were similar to those used in *in vitro* FUS stimulation to elicit calcium responses in sensory neurons^25,26^.

Figure 2 illustrates cortical responses evoked by FUS stimulation of the right CNIC. FUS elicited spiking activity in A1 (Figure 2A) in a subset of animals (n=3), whereas local field potentials (LFPs) could be recorded in most animals (n = 10, Supplementary Table 1). Representative FUS-evoked cortical LFP responses to individual stimulation triggers are shown in Figure 2B. Responses were classified as “true” when LFP negative peak amplitudes exceeded ±2 standard deviations of the pre-trigger baseline activity (dashed line). Cortical LFP responses were observed in 79% of all stimulation trials. FUS-evoked cortical responses were preceded by a brief electromagnetic artifact occurring within the first ∼8 ms following trigger onset. Across animals (n = 10), negative LFP peak latencies were generally larger than 15 ms (Figure 2C), a latency range a priori longer than that expected if afferent neurons of the CNIC were activated by FUS, provided that the process of FUS activating neurons is fast (similar to electrical excitation). Figure 2D shows average cortical LFP responses as a function of ultrasound peak positive pressure. FUS elicited cortical responses at pressures above 5 MPa, with the most robust responses observed at 6.8 MPa. Figure 2E compares cortical responses evoked by FUS and acoustic stimulation. On average, FUS-evoked cortical responses were slightly delayed relative to sound-evoked responses elicited by tone pips presented at the neuron’s best frequency and at 70 dB SPL. Furthermore, FUS evoked a broader distribution of cortical response latencies (Figure 2F). In contrast, sound-evoked responses exhibited earlier peak latencies (∼16.5 ms) and a narrower latency distribution, likely reflecting the temporally precise and synchronized responses elicited at the best frequency of neurons at the recording site (Figure 2F). Nevertheless, FUS- and sound-evoked responses exhibited similar spatiotemporal response patterns across cortical layers (Supplementary Figure 1C & D). The similarity between FUS- and sound-evoked responses suggests they share the same generation site, potentially cochlear, which is in line with previous reports of ultrasound sonication of various brain regions^17,18^. Our results also do not indicate direct FUS-induced CNIC activation. Indeed, direct activation of IC neurons by FUS would be expected to evoke substantially shorter cortical response latencies because it bypasses cochlear transduction and ascending transmission through the peripheral auditory pathway. In any case, this is what we would expect if FUS stimulation excites neurons as rapidly as electrical stimulation does^62,63^.

**Figure 2.**
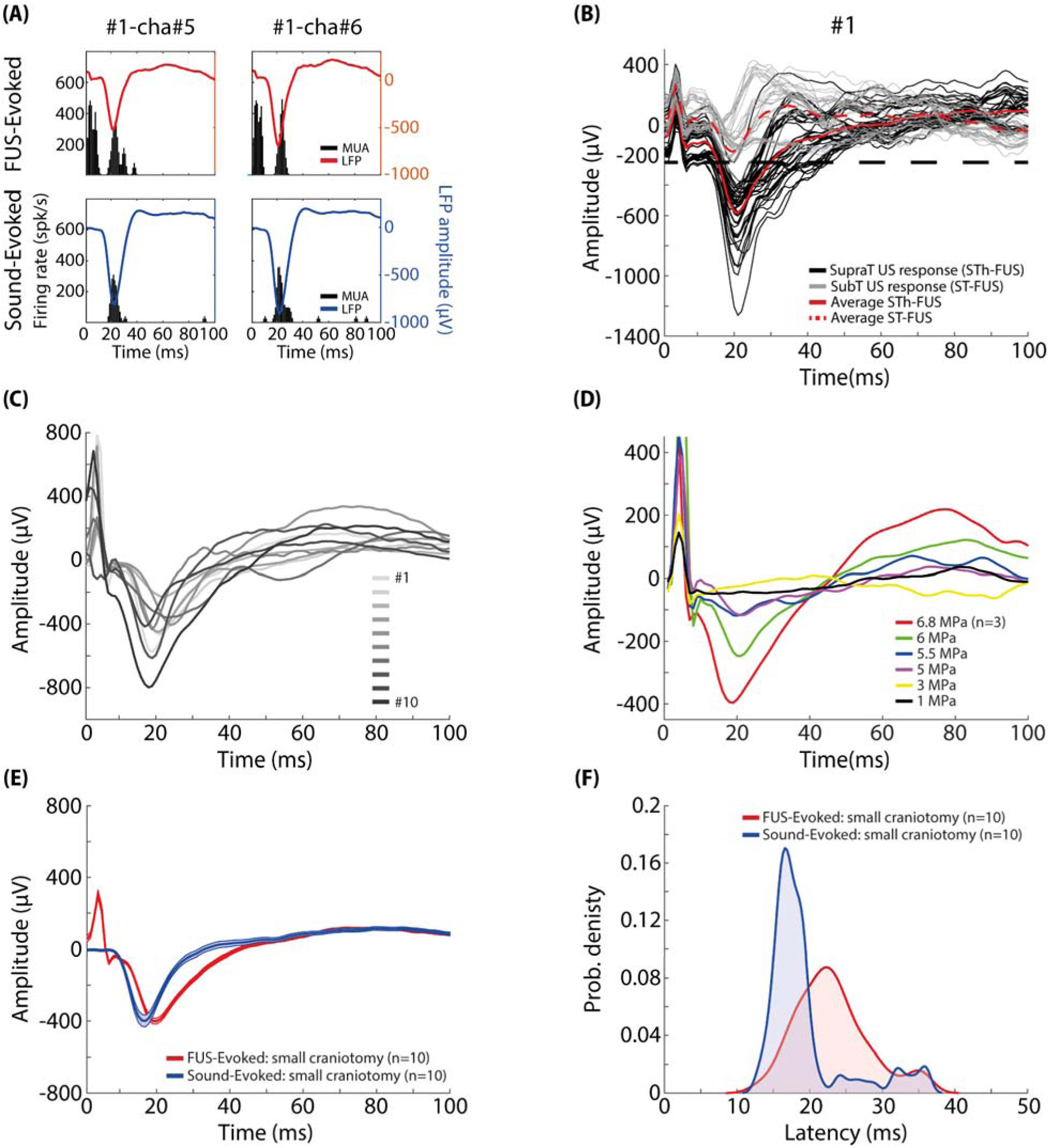
Cortical responses evoked by focused ultrasound stimulation of the inferior colliculus through small craniotomy. (A) Representative layer 4 A1 responses from two recording sites to FUS stimulation of the right inferior colliculus (IC) and acoustic stimulation (best frequency of the recording site, 70 dB SPL). For each recording site, local field potentials (LFPs) represent the average of 80 FUS trials and 10 sound trials. Post-stimulus time histograms (PSTHs; 1 ms bins) of spiking activity are shown together with the corresponding LFP traces. (B) Representative LFP responses from a layer 4 recording site across 80 FUS trials. Individual suprathreshold responses are shown in black, with the average response in red. Responses with peak amplitudes below the detection threshold (dashed line) are shown in gray, with their average indicated by the dotted red trace. (C) Averaged suprathreshold FUS-evoked LFP responses from layer 4 recording sites in each animal. (D) Averaged FUS-evoked LFP responses across ultrasound pressures. (E) Comparison of FUS-evoked (20 MHz, 6.8 MPa, 3 ms sonication duration, 90 s ISI) and sound-evoked (best frequency of the recording site, 70 dB SPL) cortical LFP responses. (F) Distribution of LFP peak latencies for FUS- and sound-evoked cortical responses.

We next wanted to confirm that the observed FUS-evoked cortical activity was, at least partly, mediated by cochlear activation. We investigated if CAPs, which reflect summed synchronous auditory nerve activity, recorded from the right (ipsilateral; n=2) and left (contralateral; n=4) cochlea could be evoked by FUS stimulation of the right IC through a small craniotomy (SC: right-FUS). Left and right columns in Figure 3 show the CAP responses in ipsi- and contralateral cochlea, respectively. In the ipsilateral cochlea (Figure 3A), FUS evoked distinct CAP responses at both ultrasound stimulus onset and offset, respectively. Brief electromagnetic artifacts (∼1 ms duration) were consistently observed at both the onset and offset of the FUS burst, preceding the CAP responses. Figure 3B illustrates the average FUS-evoked CAP response before and after death, alongside the average sound-evoked CAP (8 kHz, 70 dB SPL). Compared with the sound-evoked response, the FUS-evoked CAP exhibited a shorter N1 latency (approximately 0.1 ms), consistent with rapid cochlear activation by ultrasound. This small latency difference may be due to the additional propagation time from the earphone to the tympanic membrane and/or to the presence of an onset ramp of 1 ms for sound stimulation. Note this latency difference also contrasts with the cortical responses which showed longer latencies for FUS than for sound stimulation. No CAP responses were detected after death (Figure 3B), confirming that the responses observed in-vivo were physiological events induced by FUS rather than recording artifacts. Furthermore, FUS-evoked CAPs were detected only at peak positive pressures higher than 5 MPa (Supplementary Figure 1E), consistent with the pressure dependence of ultrasound-evoked cortical activation (Figure 2D). Unlike the ipsilateral cochlea, the contralateral cochlea did not exhibit a detectable onset FUS-evoked CAP but consistently showed an offset response in all animals (Figure 3D). This offset response was absent in post-mortem recordings (Figure 3E). Decreasing ultrasound pressure progressively reduced offset CAP amplitudes, which remained detectable down to 5.5 MPa (Supplementary Figure 1F), mirroring the pressure dependence of ipsilateral onset CAPs.

**Figure 3.**
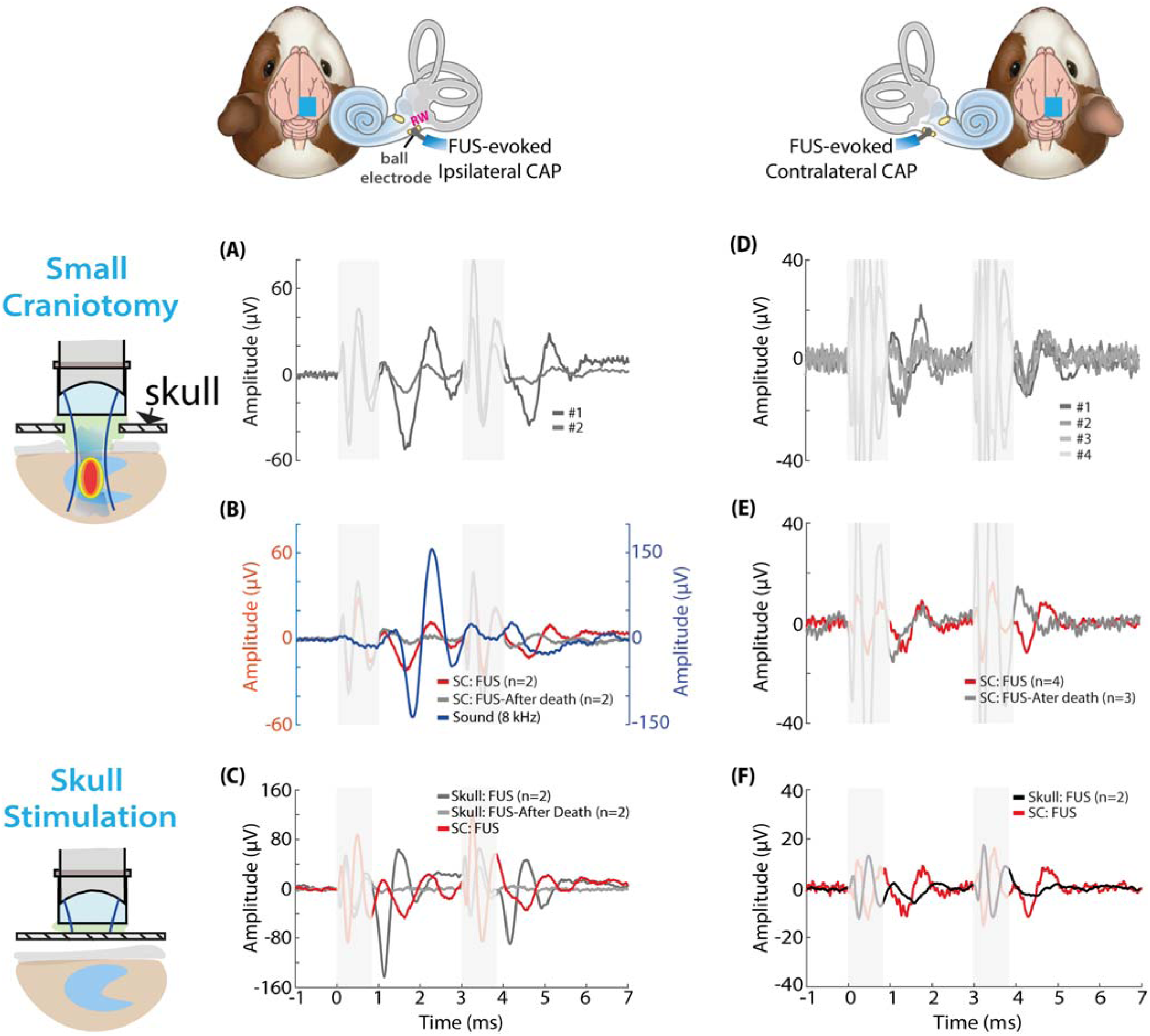
Cochlear compound action potentials evoked by FUS delivered through small-craniotomy and intact-skull stimulation. Compound action potentials (CAPs) were recorded from the ipsilateral or contralateral cochlea using a ball electrode positioned on the round window (RW) and were time-locked to each FUS trigger. (A) Representative ipsilateral CAPs evoked by FUS delivered to the IC through a small craniotomy (SC) in two animals, showing distinct onset and offset responses. Each trace represents the average of 20 FUS trials. Gray shaded regions indicate onset and offset electromagnetic artifacts. (B) Average ipsilateral FUS-evoked CAPs in the SC condition, recorded before (red) and after euthanasia (AD, gray), together with the sound-evoked CAP elicited by an 8 kHz, 70 dB SPL tone burst (blue). Line thickness indicates SEM. (C) Representative ipsilateral CAPs evoked by FUS applied over the intact skull above the IC, recorded before and after euthanasia (AD), compared with CAPs evoked by FUS delivered to the IC through a small craniotomy. (D) Representative contralateral CAPs evoked by FUS delivered to the IC through a small craniotomy in four animals, showing onset and offset responses. (E) Average contralateral FUS-evoked CAPs in the SC condition, recorded before euthanasia (red) and after euthanasia (gray). Line thickness indicates SEM. (F) Representative contralateral CAPs evoked by FUS applied over the intact skull above the IC, compared with contralateral CAPs evoked by FUS delivered to the IC through a small craniotomy.

The small craniotomy was slightly smaller than the circular aperture of the 20-MHz transducer (∼6.35 mm in diameter), allowing the focused beam to pass through the craniotomy but retaining the possibility of interaction with the surrounding skull at the craniotomy edges (Supplementary Figures 2 A1 - C1). To investigate the contribution of such skull-ultrasound interactions to FUS-induced auditory responses, we compared two additional stimulation conditions: FUS delivered through the intact skull over the right IC, and FUS delivered through a large craniotomy (LC). The large craniotomy was larger than the transducer aperture diameter, thereby minimizing direct interaction between the US focused beam and the surrounding skull (Supplementary Figures 2 A2 - C2), whereas the intact-skull condition preserved skull-ultrasound interactions. Cortical responses were recorded under both conditions. FUS stimulation of the right IC in the large craniotomy condition (n = 4, Supplementary Figures 3E & 3F) failed to evoke cortical responses at latencies comparable to those observed under SC or intact-skull conditions. In contrast, FUS stimulation through the intact skull overlying the right IC elicited robust cortical responses, with LFP onset and spike peak latencies comparable to those evoked by acoustic stimulation (8-kHz tone pips, 70 dB SPL; Supplementary Figures 3A & 3B) and closely resembling responses observed in the SC condition.

To assess cochlear involvement, FUS-evoked CAPs were recorded from the ipsilateral (n = 2) and contralateral (n = 1) cochleae in the Intact-skull condition. Robust ipsilateral CAPs with distinct onset and offset components were obtained (Figure 3C). CAP amplitudes decreased with decreasing acoustic pressure but remained detectable across all pressures tested (Supplementary Figures 3C & 3D). These responses were absent in post-mortem recordings. In contrast, contralateral CAPs exhibited only a small, longer-latency onset response, with no detectable offset component (Figure 3D). To conclude, the absence of cortical responses under the large craniotomy conditions and their persistence in the intact skull preparation point to skull-mediated mechanical coupling as a major contributor to FUS-induced cochlear activation.

### FUS stimulation recruits c-Fos activity across the central auditory pathway

In order to test if FUS can also directly stimulate IC neurons, we performed c-Fos immunohistochemistry in control, SC and LC groups at three levels of the auditory pathway: the dorsal cochlear nucleus (DCN), the inferior colliculus (IC) and the primary auditory cortex (A1). Within each structure, c-Fos-positive cells were quantified separately on the ipsilateral (FUS-stimulated) side, and the contralateral (non-stimulated) side. The contralateral side was used as an additional control (a smaller number of c-Fos positive cells are expected in the side contralateral to the FUS-stimulation, at least in the IC). In addition, we co-labeled the neurons with the NeuN marker (see Methods) to determine the proportion of neurons that express c-Fos.

The strongest and most spatially localized c-Fos response was observed in the FUS-targeted IC (Figure 4A). Whole-section mapping further revealed that c-Fos-positive cells were concentrated around the electrode trajectory and predicted ultrasound focus, with labeling increasing toward the estimated pressure maximum and decreasing outside the focal region (Figure 4A). Under both small- and large-craniotomy conditions, c-Fos expression was markedly greater in the ipsilateral than in the contralateral IC (Figures 4E & 4F). NeuN co-labeling indicated that approximately 30% of c-Fos-positive cells were neurons in the large-craniotomy condition, compared with approximately 10% in the small-craniotomy condition (Figure 4G). Together, these findings show spatially restricted cellular activation centered on the targeted IC and support focal neuronal recruitment by FUS.

**Figure 4.**
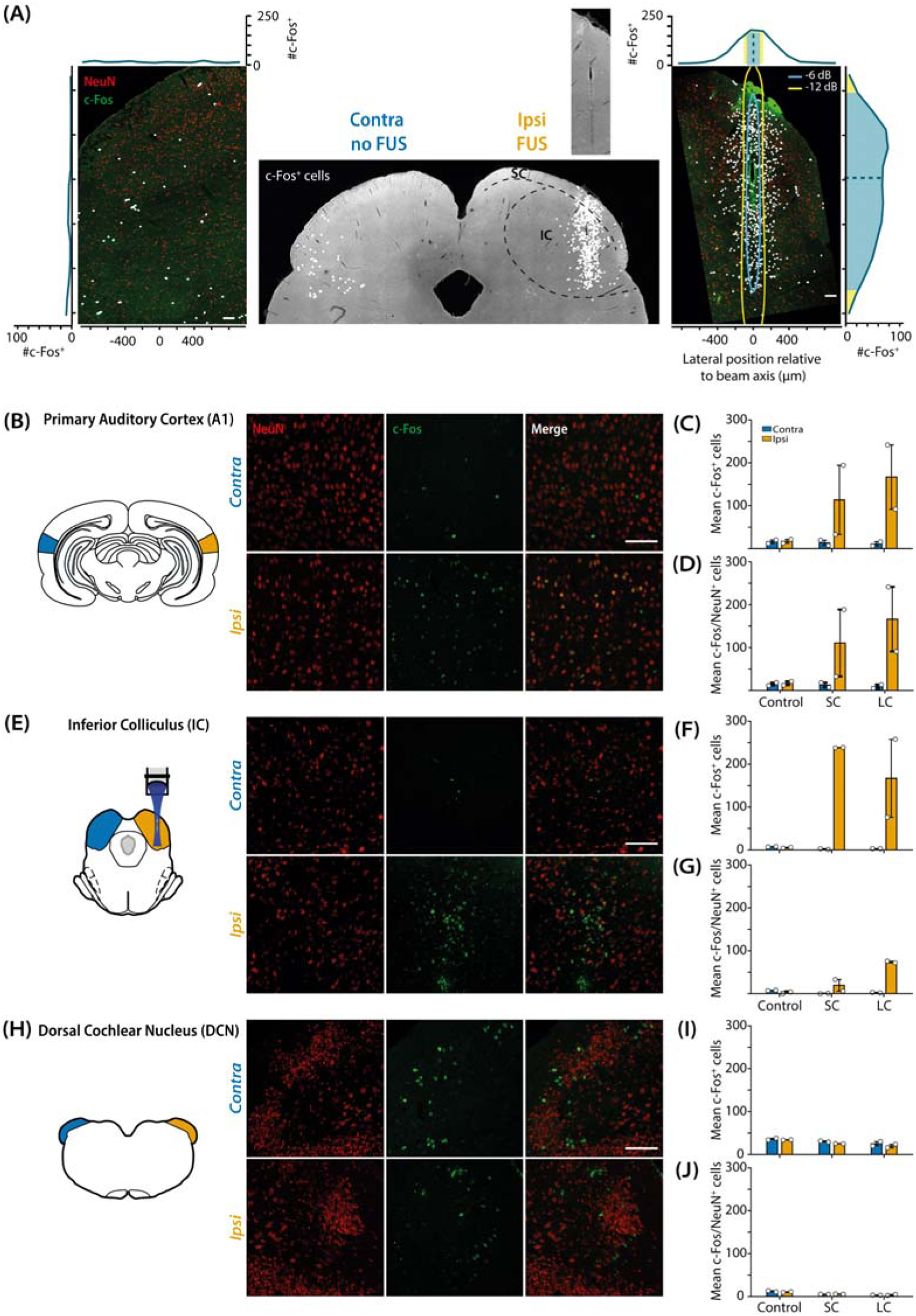
IC-targeted FUS drives region- and side-specific c-Fos activation along the auditory pathway. (A) Middle: Representative coronal section showing the spatial distribution of c-Fos-positive cells (white) in the inferior colliculus (IC) following FUS stimulation of the ipsilateral IC. Inset (top) shows the electrode trace in the IC, used to confirm targeting accuracy. The approximate border of the IC and superior colliculus (SC) are indicated for reference (dotted curved line). Left and right: Enlarged regions of interest from the contralateral and ipsilateral IC, with NeuN (red) and c-Fos (green) labelling. c-Fos-positive cells are additionally indicated by white dots for visualization. In the ipsilateral IC, c-Fos-positive cells were concentrated along the acoustic beam axis, predominantly within the simulated −6- (blue) and −12-dB (yellow) pressure contours. Side plots show the spatial distribution of c-Fos positive cells counts along the dorsoventral axis (left and right plots) and the mediolateral axis (top plot) for both the ipsilateral and contralateral IC. (B,E,H) Representative NeuN, c-Fos, and merged images from contralateral and ipsilateral primary auditory cortex (A1), inferior colliculus (IC), and Dorsal Cochlear Nucleus (DCN). (C,F,I) Quantification of c-Fos-positive cells across the three structures under control, small-craniotomy (SC), and large-craniotomy (LC) conditions. (D,G,J) Quantification of c-Fos/NeuN double-positive cells across the three structures under control, small-craniotomy (SC), and large-craniotomy (LC) conditions. Scale bar = 100µm

FUS-induced c-Fos expression was also detected in A1 under both small- and large-craniotomy conditions (Figures 4B - 4C) where c-Fos-positive cells were quantified in layers IV and V/VI (see Supplementary Figure 4A & 4B for spatial distribution of c-Fos activation across cortical layers). In the small-craniotomy condition, fewer c-Fos-positive cells were detected in ipsilateral A1 (113 ± 80) than in ipsilateral IC (238 ± 1), whereas contralateral A1 and IC counts were both comparatively low (13.5 ± 6 and 2 ± 1, respectively). In the large-craniotomy condition, ipsilateral A1 and IC showed similar c-Fos counts (167 ± 75 and 167 ± 91, respectively). NeuN co-labeling showed that nearly all c-Fos-positive cells in A1 were neuronal in both the small- and large-craniotomy conditions (∼98% in each FUS group, Figure 4D).

In contrast, c-Fos labeling in the dorsal cochlear nucleus (DCN; Figures 4H – J) remained relatively low across control (ipsilateral: 34 ± 1; contralateral: 36 ± 2), small-craniotomy (25 ± 1; 30 ± 1), and large-craniotomy conditions (20 ± 4; 25 ± 5). Most c-Fos-positive cells were localized to the granule-cell region of the DCN (Supplementary Figures 4C - F), whereas little or no c-Fos expression was detected in the ventral cochlear nucleus (Supplementary Figures 4G & 4H).

In summary, FUS produced spatially restricted c-Fos activation centered on the targeted IC, with NeuN co-labeling confirming neuronal recruitment within the stimulated region. c-Fos expression in A1 further suggests engagement of the auditory afferent pathway.

## Sustained effects of FUS-induced cochlear activation

Our results so far show that FUS induces excitation of the auditory afferent pathway in two different ways including indirect cochlear excitation and direct IC activation. We next asked whether FUS induces lasting changes at the cochlear level, either as a consequence of the multiple cochlear activation indirectly induced by FUS stimulation or through direct activation of the auditory efferent system^64,65^ induced by FUS at the level of the CNIC. Therefore, to determine whether FUS stimulation of the right IC can modulate sound-evoked cochlear responses, CAPs and cochlear microphonics (CMs) were recorded from the contralateral cochlea in response to 4- and 8-kHz tone pips (alternating polarity, 60 dB SPL) presented to the same ear. Measurements were obtained before, during, immediately after, and up to 1 h following the 2-h FUS stimulation period. Figures 5A & 5B show representative CM and CAP responses recorded before, +60 min after FUS, and after FUS stimulation sequence under SC and LC conditions. All averaged data are shown in Figures 5C - 5F. CM and CAP amplitudes at both 4 and 8 kHz remained stable across baseline recordings obtained 1h apart before FUS stimulation, indicating stable cochlear recordings in the absence of ultrasound stimulation.

**Figure 5.**
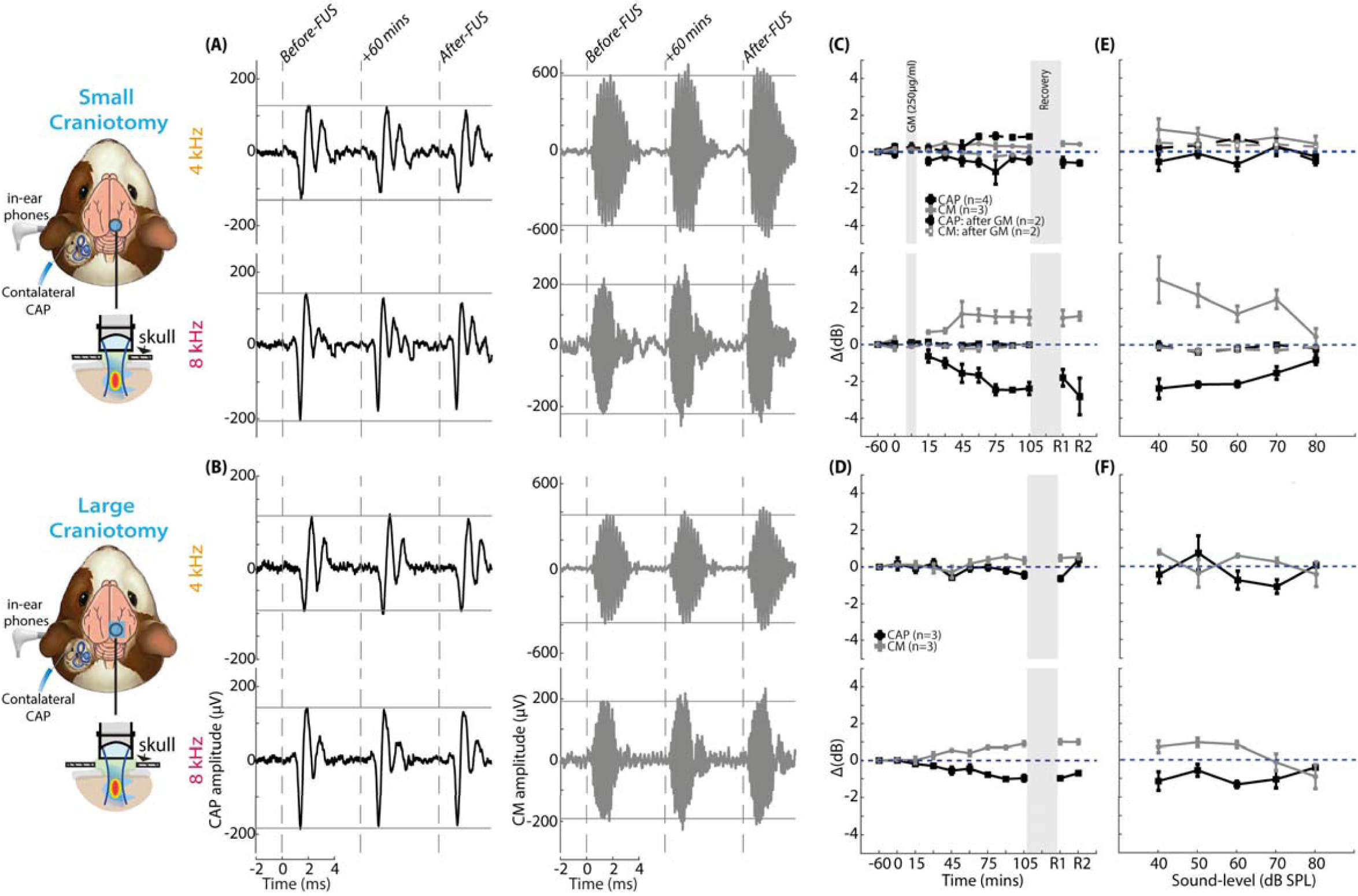
IC-targeted FUS engages auditory efferent pathways and produces sustained modulation of cochlear responses. (A,B) Left: Schematic of the Experimental paradigms for recording contralateral sound-evoked compound action potentials (CAPs) and cochlear microphonics (CMs) during FUS stimulation of the right inferior colliculus through a small (A) or large (B) craniotomy are shown. Representative CAP and CM responses evoked by 4- and 8-kHz tone bursts (60 dB SPL) are shown at baseline, at the end of the stimulation period and 60 min after the FUS stimulation sequence. (C) Time course of normalized CAP (black) and CM (gray) amplitudes evoked at 4 and 8 kHz (60 dB SPL) during FUS delivered through a small craniotomy. In two animals, the contribution of the auditory efferent pathway was assessed by measuring responses before gentamicin (GM) administration, 3 h after GM administration immediately before FUS, and throughout the FUS stimulation period. Dotted lines indicate the mean CAP and CM amplitudes measured after GM treatment and before FUS onset. (D) Time course of normalized CAP (black) and CM (gray) amplitudes evoked at 4 and 8 kHz (60 dB SPL) before, during, and after FUS delivered through a large craniotomy. (E) Normalized CAP and CM amplitudes as a function of sound level (40–80 dB SPL) following FUS delivered through a small craniotomy, shown under intact efferent conditions and after gentamicin treatment. (F) Normalized CAP and CM amplitudes as a function of sound level (40–80 dB SPL) following FUS delivered through a large craniotomy.

Small changes in CM and CAP amplitudes were observed at 4 kHz throughout the 2-h stimulation sequence and after FUS in both stimulation conditions (SC and LC) (Figures 5C & 5D). In contrast, at 8 kHz, FUS through the SC induced numerically larger changes in CM and CAP amplitudes, detectable within 1 5 min of FUS onset (CM: 0. 67 ± 0. 06 dB; CAP: −0.61 ± 0. 3 dB). These effects increased progressively, reaching a plateau by 90 min of stimulation (CM: 1 . 5 ± 0. 32 dB; CAP: −2. 4 ± 0. 27 dB) that persisted for the remainder of the stimulation period. Notably, CM and CAP amplitudes remained altered throughout recovery and failed to return to baseline 1 h after FUS under both stimulation conditions. In the LC condition at 8 kHz, FUS stimulation induced a progressive increase in CM amplitude and decrease in CAP amplitude, emerging 45 min after FUS onset (CM: 0. 54 ± 0. 04 dB; CAP: −0. 56 ± 0. 1 9 dB) and reaching 0. 91 ± 0. 11 dB and −0. 97 ± 0. 1 5 dB, respectively, by the end of stimulation. These observations, namely the enhancement of CM amplitude and suppression in CAP amplitude, particularly prominent at 8 kHz, are consistent with an activation of the MOC system by FUS stimulation targeting the CNIC^66, 67^.

To confirm FUS-induced changes in CM and CAP amplitudes were mediated by efferent activation, gentamicin (GM; 250 mg/kg, n = 2) was administered intraperitoneally 2 h before FUS stimulation. This dose was chosen based on previous studies in guinea pigs showing that it can attenuate cholinergic efferent transmission at the level of outer hair cells without inducing immediate structural damage to the cochlea^68^. CM and CAP responses were recorded before GM administration, 2 h after injection just before FUS stimulation sequence, throughout the 2-h FUS stimulation sequence, and immediately after FUS stimulation sequence (Figures 5C-5E). Following GM treatment, FUS no longer altered CM or CAP amplitudes at 8 kHz. At 4 kHz, however, GM reversed the FUS effect on CAP amplitudes, producing a selective increase in CAP responses without affecting CM amplitudes (Supplementary Figure 5). These findings indicate that the FUS-induced modulation of CM and CAP responses observed under control conditions in the absence of GM is mediated by the activation of the MOC system.

The level dependence of FUS-induced cochlear modulation was assessed by measuring CM and CAP amplitudes at 4 and 8 kHz before and after FUS stimulation of the right IC through SC and LC (Figures 5E & 5F). At 4 kHz, FUS-induced changes were modest across stimulus intensities. In contrast, at 8 kHz, CM and CAP changes were largest at lower sound levels, particularly in the SC condition (Figure 5E), and diminished at higher intensities.

A generalized linear mixed-effects model (GLMM; Gamma distribution with log-link function; see Methods) was used to evaluate FUS-associated changes in CM and CAP amplitudes. CM amplitudes showed significant main effects of Time (χ² = 6.55, p = 0.0105), Group (SC and LC) (χ² = 10.29, p = 0.0058), Frequency (χ² = 27.54, p = 1.54 × 10⁻O), and Intensity (χ² = 398.12, p < 0.001). CAP amplitudes similarly showed significant main effects of Time (χ² = 7.53, p = 0.0061), Group (χ² = 4.76, p = 0.029), Frequency (χ² = 35.86, p = 2.12 × 10⁻O), and Intensity (χ² = 110.37, p < 0.001). Because of the relatively small group sizes, interaction terms were not retained in the primary models (Supplementary Table 2). The significant effect of Group reflects overall differences in CM and CAP amplitudes between the LC and SC groups. However, this effect does not imply that the FUS-induced change in CAP or CM amplitude differed between SC and LC: complementary analyses testing specifically whether the Pre-to-Post changes differed between groups revealed that the addition of a Time × Group interaction did not improve model fit for either CM or CAP.

## The protective effect of FUS against noise trauma

We described above that FUS stimulation induces cortical responses similar to those induced by acoustic stimulation. Very soon after this first result, we suspected that these responses were linked to cochlear activation^17,18^. To test this hypothesis quickly and simply, we then exposed the animals to an acoustic trauma (continuous 8 kHz-tone at 115 dB SPL for 1 hour) immediately after the 2-hours FUS stimulation sequence. We hypothesized that if acoustic trauma-induced hearing loss modified or abolished the FUS-related cortical responses, it would support the idea that these responses are linked to direct cochlear activation by FUS stimulation. As the experiment progressed, we noticed that the animals previously exposed to ultrasound and then to acoustic trauma showed little or no hearing loss. We therefore documented this unexpected effect in 6 animals. A schematic of the experimental setup is shown in Figure 6A.

**Figure 6.**
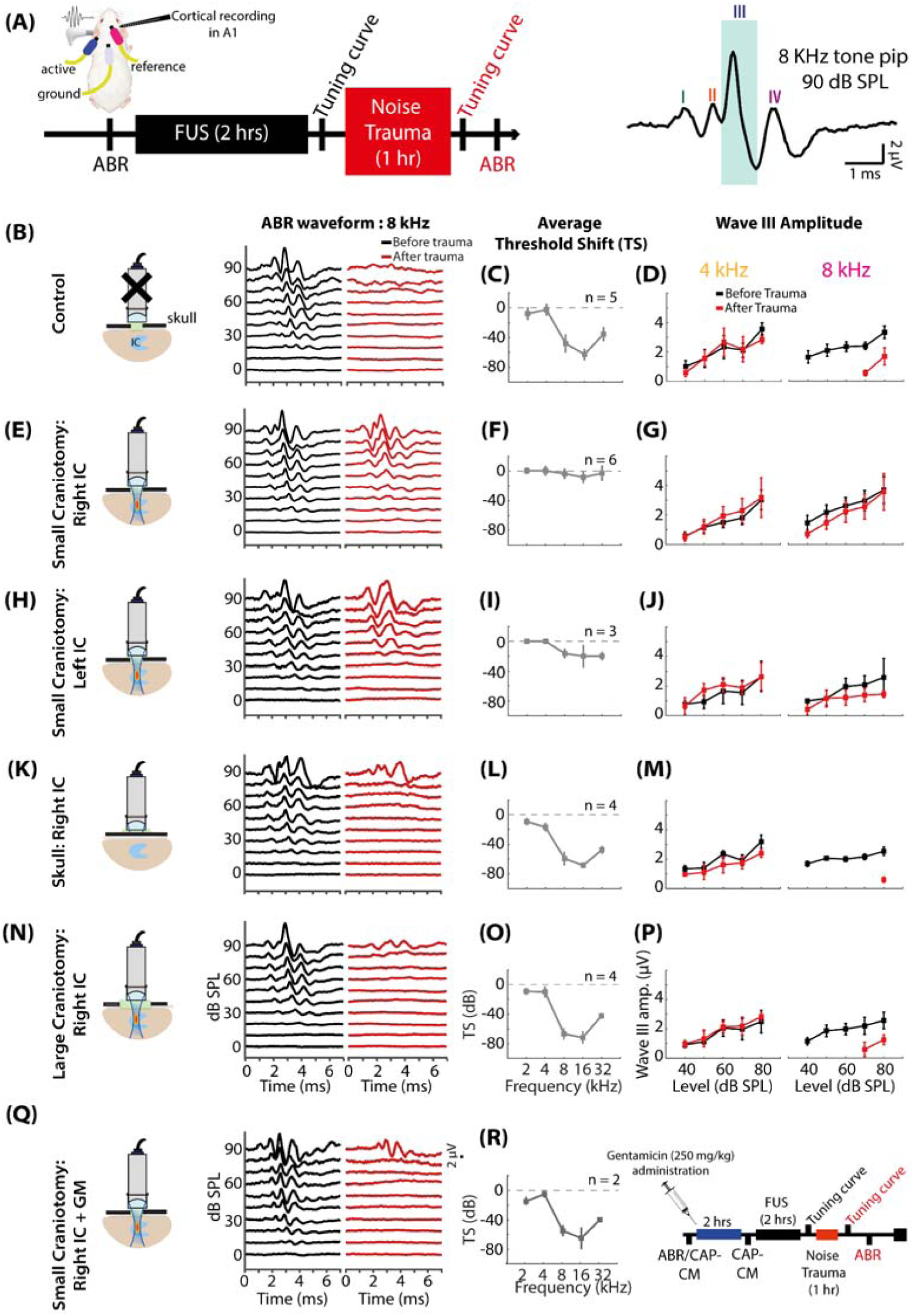
FUS stimulation of the inferior colliculus protects against noise-induced hearing loss. (A) Experimental paradigm used to assess the effect of IC-targeted FUS on noise-induced hearing loss. ABRs were recorded before and after acoustic trauma, and sound-evoked cortical responses were recorded from A1 before and after noise exposure. Right, representative ABR waveform evoked by an 8-kHz tone pip at 90 dB SPL, showing waves I–IV; wave III, used for threshold determination, is highlighted. (B–P) from left to right columns: Representative 8-kHz ABR waveforms, mean ABR threshold shifts across frequencies, and wave III amplitude-level functions at 4 and 8 kHz before and after acoustic trauma for Control (B–D**; n = 5)**, small-craniotomy right-IC FUS (E–G**; n = 6)**, small-craniotomy left-IC FUS (H–J**; n = 3)**, intact-skull right-IC FUS (K–M**; n = 4)**, and large-craniotomy right-IC FUS (N–P**; n = 4)** conditions. (Q and R) Effect of auditory efferent blockade on FUS-mediated protection. Gentamicin (GM; 250 mg/kg) was administered before right-IC FUS through a small craniotomy. (Q) Representative 8-kHz ABR waveforms before and after acoustic trauma. (R) Mean ABR threshold shifts across frequencies (n = 2) and schematic of the experimental sequence. Data are shown as mean ± SEM. ABR, auditory brainstem response; A1, primary auditory cortex; FUS, focused ultrasound; IC, inferior colliculus; GM, gentamicin.

We also added a control group, where animals (n = 5) underwent the same surgical preparation and experimental setup, but for which no FUS stimulation was delivered. Animals were instead maintained in silence for 2 h before undergoing the same noise-exposure protocol. Auditory brainstem responses (ABRs) were recorded from the left ear before and after noise exposure to assess Noise-Induced Hearing Loss (NIHL). Thresholds were assessed using wave III^39^ (Figure 6A). In control animals, acute noise exposure induced a 60-dB hearing loss at and above 8 kHz (Figures 6B & 6C), consistent with our previous studies using the same noise exposure paradigm^41,69^. It should be noted that the “smaller” threshold shift at 32 kHz is an “artefact” resulting from the fact that the maximum level used to collect the ABRs was 90 dB SPL. In contrast, animals exposed to FUS stimulation on the right IC (Figure 6E & 6F) exhibited ABR threshold elevations of less than 10 dB at and above 8 kHz following noise exposure, indicating a substantial protective effect of FUS against NIHL. Individual ABR threshold shifts for all animals and experimental conditions are shown in Supplementary Figures 6B - 6G. Next, we investigated whether stimulating the left IC (Figures 6H & 6I) with FUS could also have a protective effect against noise trauma. This condition also provided substantial protection (approximately 20 dB), albeit slightly less than that provided by right IC stimulation. Protection against the effects of acoustic trauma is approximately 50 dB for acute hearing loss—that is, the hearing loss that occurs immediately after acoustic trauma. For comparison, we wanted to determine the extent of chronic hearing loss two weeks after the same acoustic trauma in a separate group of animals (n=3), namely, once hearing loss had stabilized following spontaneous recovery of hearing. Chronic hearing loss is approximately 30 dB at 8 and 16 kHz (Supplementary Figure 6A).

We next wanted to study whether cochlear and/or possibly efferent activation by FUS stimulation was sufficient to confer protection against acoustic trauma. To this end, we compared NIHL across two additional FUS paradigms: stimulation through the intact skull (Skull condition, Figures 6K & 6L) and stimulation of the right IC through a large craniotomy (LC: right-FUS, Figures 6N & 6O). Based on our previous experiments (see above), it was expected that these conditions would engage cochlear activation only, and CNIC stimulation (including excitation of efferent neurons) with minimal cochlear activation, respectively. Strikingly, protection against acoustic trauma was absent in both the Skull and LC conditions, both groups exhibiting high-frequency threshold shifts comparable to controls (∼60 dB above 8 kHz; Figures 6B & 6C).

A linear mixed-effects model revealed significant main effects of Time (F(1,153) = 275.26, p < 0.001) and Frequency (F(4,153) = 137.50, p < 0.001), whereas the main effect of Group was not significant (F(4,17) = 2.11, p = 0.124). Significant Time × Group (F(4,153) = 29.01, p < 0.001), Time × Frequency (F(4,153) = 31.05, p < 0.001), Group × Frequency (F(16,153) = 8.29, p < 0.001), and Time × Group × Frequency interactions (F(16,153) = 2.88, p = 3.90 × 10⁻O) were observed, indicating that the effect of noise exposure differed across experimental groups and frequencies. Given the significant three-way interaction, Bonferroni-adjusted interaction contrasts were used to directly compare the pre-to-post threshold changes between groups at each frequency. Noise-induced threshold shifts were significantly smaller in the SC:right-FUS group than in the Control, LC, and Skull groups at 8 kHz (adjusted p = 1.77 × 10⁻O, 9.56 × 10⁻O, and 4.03 × 10⁻O, respectively), 16 kHz (adjusted p = 1.28 × 10⁻O, 7.71 × 10⁻O, and 4.60 × 10⁻O), and 32 kHz (adjusted p = 0.0025, 4.59 × 10⁻O, and 5.65 × 10⁻O), whereas no significant differences were detected at 2 or 4 kHz. Threshold shifts in the SC:right-FUS group did not significantly differ from those in the SC:left-FUS group at any tested frequency (all adjusted p > 0.47). Consistent with these between-group comparisons, the mean pre-to-post threshold change across frequencies was minimal in the SC:right-FUS group (+2.7 dB; p = 0.378), compared with larger increases in the Control (+31.3 dB), LC (+40.0 dB), and Skull (+40.1 dB) groups.

We next assessed changes in ABR wave III amplitudes across experimental groups (Figures 6D, G, J, M, P) following noise exposure. A generalized linear mixed-effects model (Gamma distribution with log-link function) revealed significant effects of Time (χ² = 47. 84, p < 0. 001), Frequency (χ² = 8. 44, p = 0. 0037), and Intensity (χ² = 1 5. 57, p < 0. 001), whereas the main effect of Group was not significant (χ² = 4. 75, p = 0. 1 91). Significant Time × Group (χ² = 1 3. 1 9, p = 0. 0043), Time × Frequency (χ² = 9. 02, p = 0. 0027), and Group × Frequency (χ² = 11 . 32, p = 0. 01 0) interactions were observed. Bonferroni-adjusted contrasts directly comparing the pre-to-post changes between groups showed that the reduction in wave III amplitude was significantly smaller in the SC group than in the Control (z = −2. 42, adjusted p = 0. 047), LC (z = −2. 84, adjusted p = 0. 01 3), and Skull (z = −3. 33, adjusted p = 0.0026) groups.

Finally, we tested on two animals whether the protection against noise exposure observed in the SC:right-FUS group was mediated by FUS-induced activation of the efferent auditory system. To this end, gentamicin (GM; 250 mg/kg) was administered intraperitoneally before FUS stimulation sequence (Figure 6R, right). Following GM administration, noise exposure after FUS resulted in a ∼60-dB ABR threshold shifts at and above 8 kHz (Figures 6Q & 6R). These findings indicate that the protective effect of FUS depends on the activation of the efferent auditory system together with cochlear activation.

## Cortical changes after noise trauma

Cortical consequences of acute noise exposure were assessed from tone-evoked LFP tuning curves (0.5–32 kHz, 1/8-octave steps; 20–70 dB SPL). Mean ABR threshold shifts for each experimental condition are replotted in Figures 7A – K to facilitate comparison between peripheral hearing loss and cortical responses. Layer IV LFP tuning curves were aligned to the characteristic frequency (CF) of each recording site and averaged across sites, with frequency expressed in octaves relative to CF (Figures 7B – L). Representative pre- and post-noise Layer IV LFP tuning curves from a single recording site for each experimental condition are shown in Supplementary Figures 6B – 6G. Cluster-based permutation statistical maps highlighting significant cortical changes are also shown (at the right side of cortical responses). Noise exposure significantly reduced cortical responses in Control (Figure 7B), intact-skull: right-FUS (Figure 7H), and LC: right-FUS animals (Figure 7J; p < 0.05), consistent with previous observations following acute noise exposure^51^. In contrast, cortical responses remained largely preserved in both SC: right-FUS (Figure 7D) and SC: left-FUS groups (Figure 7F). Consistent with this preservation, current-source density (CSD) profiles following SC: right-FUS showed no apparent changes in the spatiotemporal organization of cortical sinks and sources after noise exposure compared with pre-noise responses (Supplementary Figures 7A & 7B). In addition, some responses were unmasked at specific frequency–intensity combinations (low-frequency, 70 dB SPL), particularly in the SC: right-FUS group. Consistent with the loss of ABR protection following gentamicin treatment, preservation of cortical tuning was also abolished in the SC: right-FUS group condition, with cortical responses reduced following noise exposure (Figure 7L).

**Figure 7.**
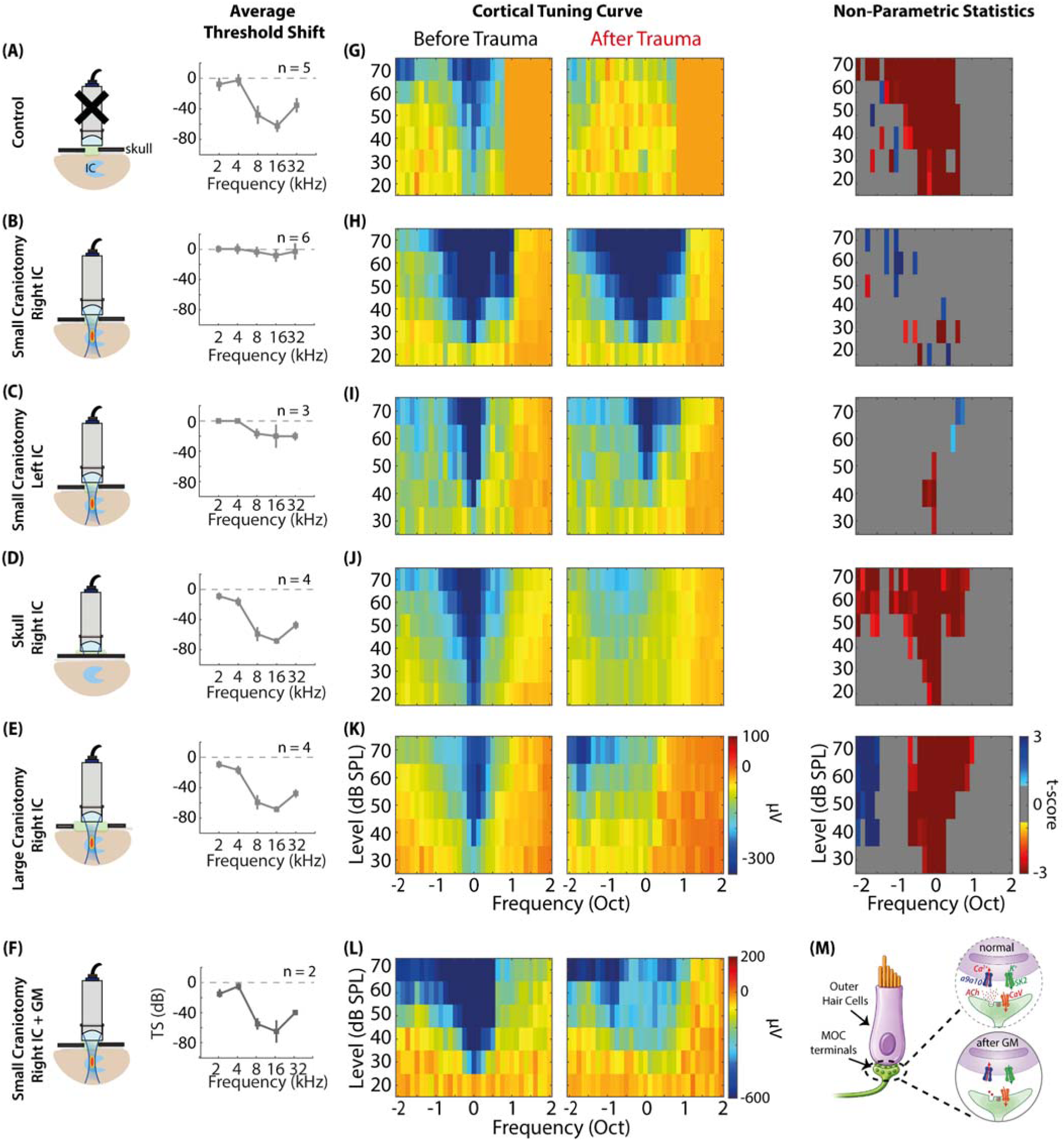
IC-targeted FUS preserves auditory cortical tuning following acoustic trauma under protective stimulation conditions. (A–K) Experimental conditions and corresponding mean ABR threshold shifts following acoustic trauma for Control (no FUS; n = 5), small-craniotomy right-IC FUS (n = 6), small-craniotomy left-IC FUS (n = 3), intact-skull right-IC FUS (n = 4), large-craniotomy right-IC FUS (n = 4), and small-craniotomy right-IC FUS following gentamicin treatment (GM; n = 2), respectively. Mean ABR threshold shifts are replotted from Figure 7 as a reference for the degree of peripheral hearing loss associated with each condition. (B–L) Corresponding averaged auditory cortical LFP tuning maps before and after acoustic trauma. Frequency is expressed in octaves relative to the characteristic frequency of each recording site. Statistical maps (on the right) show significant pre- versus post-trauma differences identified using cluster-based non-parametric permutation testing; red and blue indicate significant decreases and increases in response magnitude, respectively, whereas gray indicates non-significant differences. (M) Schematic of the medial olivocochlear (MOC) efferent synapse onto outer hair cells, illustrating normal cholinergic efferent signaling and its disruption following gentamicin treatment. ACh, acetylcholine; ABR, auditory brainstem response; FUS, focused ultrasound; GM, gentamicin; IC, inferior colliculus; LFP, local field potential; MOC, medial olivocochlear; TS, threshold shift.

### Safety Characterization of FUS stimulation and simulations

We next asked whether FUS stimulation parameters used in this study induced damage to target IC tissues. To examine cellular integrity and apoptosis following FUS stimulation (6.8 MPa, 3-ms pulses, 90-s interstimulus interval), regions of the right-IC within the FUS focus were analyzed using DAPI and caspase-3. DAPI staining showed preserved nucleus organization, and cleaved caspase-3 labeling was not increased in the stimulated IC relative to controls (Figures 8A & 8B). These data suggest that the FUS parameters employed in this study do not induce apoptotic damage to targeted IC tissues.

**Figure 8.**
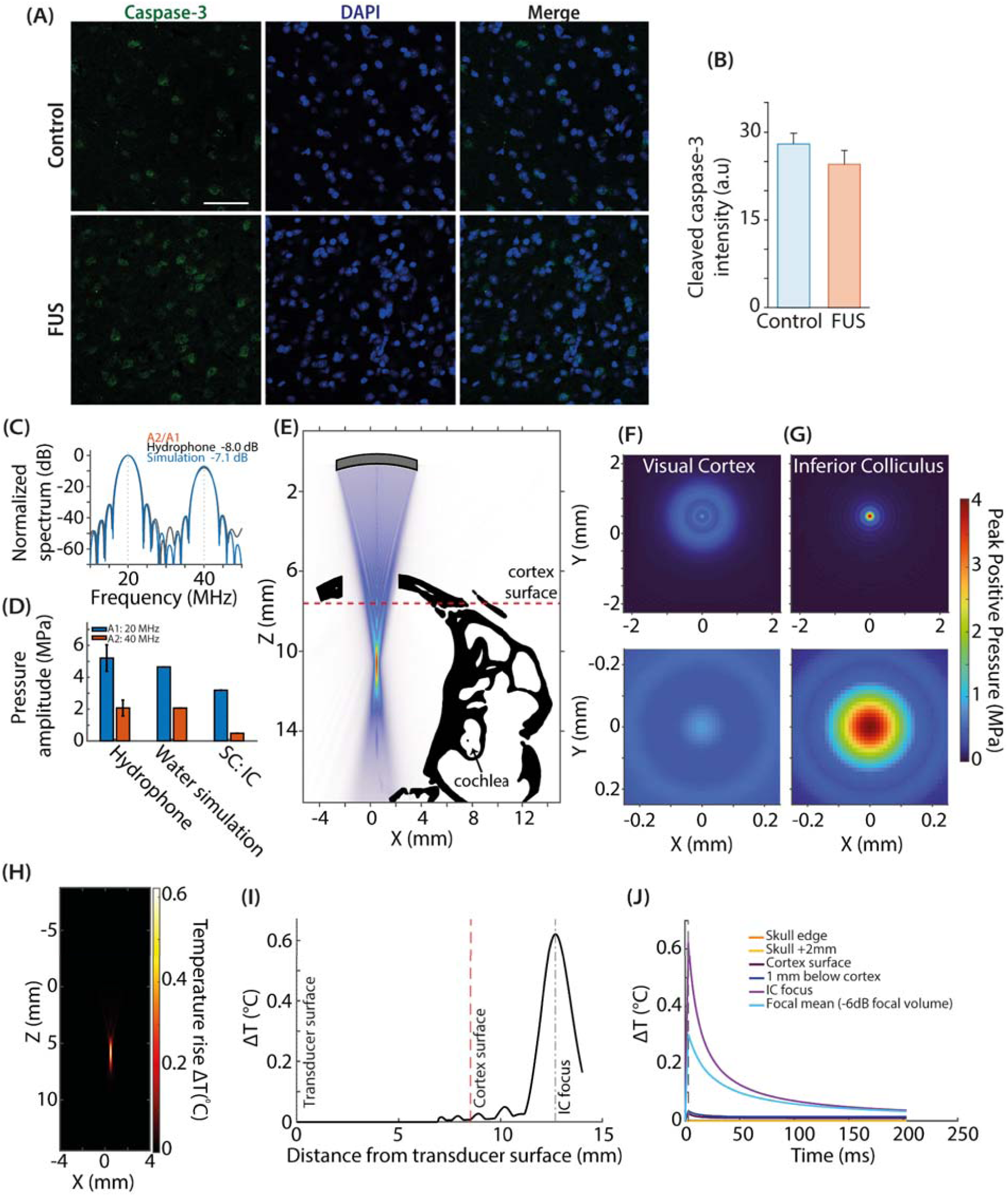
Cell death and 3D acoustic and thermal characterization of FUS exposure at the inferior colliculus. (A) Representative cleaved caspase-3 (green), DAPI (blue), and merged images from control and FUS-stimulated IC (Scale bar = 50 µm). (B) Quantification of cleaved caspase-3 fluorescence intensity in control and FUS-stimulated IC. (C) Normalized pressure spectra measured experimentally with the hydrophone and obtained from the corresponding water simulation. Dashed lines indicate the 20-MHz fundamental (A1) and 40-MHz second harmonic (A2); A2/A1 ratios are indicated. (D) A1 and A2 pressure amplitudes measured with the hydrophone and estimated from the water and small-craniotomy IC (SC:Right-IC) simulations. Hydrophone values are shown with calibration uncertainty. (E) Simulated acoustic field overlaid on the CT-derived anatomy (small craniotomy), showing the axial peak-positive pressure field. The dashed red line indicates the cortical surface. (F-G) Transverse pressure distributions 1 mm below the visual cortical surface and at the IC focal plane (top). Enlarged transverse pressure maps at the corresponding cortical and IC focal planes, shown in bottom. (H) Spatial distribution of FUS-induced temperature rise through the small craniotomy. (I) Simulated temperature rise along the acoustic propagation axis from the transducer surface to the IC focus. Dashed lines indicate the cortical surface and IC focal position. (J) Temporal evolution of temperature rise at selected locations along the propagation path and averaged across the focal region following a 3-ms FUS pulse. Bars show mean ± SEM. A1, 20-MHz fundamental; A2, 40-MHz second harmonic; IC, inferior colliculus; SC, small craniotomy; FUS, focused ultrasound.

To estimate the acoustic pressure reaching the IC and the associated thermal effects, we performed three-dimensional nonlinear acoustic simulations using the k-space pseudospectral method implemented in k-Wave^54^. The simulations were first calibrated by adjusting the transducer excitation pressure to match the amplitudes of the 20-MHz fundamental (A1) and 40-MHz second-harmonic (A2) components measured with a hydrophone at the beam focus in water (see Methods and Figures 8C & 8D).

The simulated amplitudes of the 20-MHz fundamental (A1 = 4. 7 MPa) and 40-MHz second harmonic (A2 = 2. 05 MPa) yielded an A2/A1 ratio of −7. 1 dB, consistent with the hydrophone measurements which yielded an A2/A1 ratio of -8 dB (A1 = 5. 2 ± 0. 8 MPa and A2 = 2. 1 ± 0. 5 MPa, where the ± values represent the measurement uncertainties specified by the needle hydrophone manufacturer) (Figure 8D). We then simulated nonlinear ultrasound propagation through the brain, with the beam focused on the IC, while accounting for the effects of the skull with small craniotomy, using a 3D anatomical model based on yCT scan (Figure 8E). Transversal pressure maps obtained 1 mm below the cortical surface and at the IC focal plane showed progressive spatial concentration of the beam toward the target, with a confined pressure maximum within the IC (Figures 8F and 8G). After propagation through the intermediate attenuating tissues, the simulated pressures at the IC were approximately 3.1 MPa at 20 MHz and 0.4 MPa at 40 MHz, corresponding to ∼66% and to ∼20% of the respective free-field simulated amplitudes (4.7 MPa at 20 MHz and 2.05 at 40 MHz; Figure 8D). Thus, propagation through the heterogeneous anatomical path attenuated the acoustic pressure at the IC, with a stronger reduction of the 40-MHz second-harmonic component than of the 20-MHz fundamental, while preserving spatial focusing at the IC. We finally quantified FUS-induced heating in brain tissue by solving the bioheat equation. The temperature rise was localized near the IC focus, reaching a maximum of approximately 0.6–0.7°C, while remaining below 0.1°C at the cortical surface and along the US propagation path through the brain (Figures 8H and 8I). At the IC, temperature peaked at the end of the 3-ms sonication and rapidly declined over the following few hundred milliseconds (Figure 8J). The estimated peak temperature rise at the IC remained below 0.8°C, below the 2°C threshold below which the thermal risks are considered nonsignificant^70^.

## Discussion

### FUS activates the cochlea

Our study revealed that our FUS stimulation targeting the CNIC via a small craniotomy can activate the auditory cortex, likely through indirect activation of the cochlea. These results are consistent with the literature^17,18^ and are supported by several pieces of evidence. First, we show that FUS stimulation is associated with CAP responses in the ipsilateral ear and to a smaller extent with CAP responses in the contralateral ear. This indirect cochlear activation is likely causing cortical responses which were also observed when FUS stimulation was applied directly to the intact skull. At the high frequency used in this study (20 MHz), FUS is unable to penetrate the intact skull and cannot, therefore, modulate neural activity in the CNIC. Second, the latencies of the cortical responses (>20 ms) also appear more consistent with cochlear-induced activation of A1 than with direct activation of afferent neurons in the CNIC. To further investigate the mechanisms underlying FUS-induced cochlear activation, we introduced an additional condition involving a large craniotomy (LC) to minimize interactions between the focused ultrasound beam and the surrounding skull while retaining sonication of the CNIC (Supplementary Figures 2 A2 - C2). FUS stimulation under this condition failed to induce either cochlear or cortical responses (Supplementary Figures 3E & 3F), suggesting that the FUS-evoked cortical responses reflect activity originating from the cochlea.

Several mechanisms could potentially explain cochlear activation following FUS through the small craniotomy, including mechanical interactions with the skull and/or cerebrospinal fluid (CSF)-mediated effects. The ultrasonic field can interact with the surrounding skull at the craniotomy edges (Supplementary Figures 2 A1 – C1), and previous work at lower ultrasound frequencies (200 kHz) has proposed that the skull may act as a waveguide for shear waves propagating toward off-target structures such as the cochlea^22^. Alternatively, FUS may generate mechanical forces through acoustic radiation force and acoustic streaming, producing fluid motion in the CSF within the subarachnoid space exposed by the craniotomy^71^ and potentially transmitting mechanical perturbations toward the cochlea through the cochlear aqueduct. Our experimental design does not allow us to distinguish between these two mechanisms. In the intact skull condition, the strong attenuation and limited transmission of 20-MHz ultrasound through bone would substantially reduce the acoustic pressure reaching the CSF within the subarachnoid space, making a direct CSF-mediated mechanism less likely under these conditions. Thus, the persistence of cochlear and cortical responses with the intact skull, together with their absence following large-craniotomy stimulation, is consistent with a contribution of skull-mediated mechanical coupling even if a contribution from CSF-mediated effects cannot be excluded.

### FUS targeting CNIC activates the afferent and efferent pathways

Notably, c-Fos-positive cells were consistently observed in the IC and auditory cortex ipsilateral to FUS stimulation under the large-craniotomy condition, in which cochlear activation was minimized. In contrast, contralateral labeling remained sparse and comparable to control levels, consistent with background activity rather than FUS-driven recruitment. Within the IC, c-Fos-positive cells were concentrated around the targeted region and largely confined to the −6 dB contour of the FUS beam, supporting spatially localized recruitment by the 20-MHz stimulus (Figure 5A). Because c-Fos is an indirect marker of recent cellular activation that does not directly report neuronal firing and can also be expressed in non-neuronal cells, neuronal recruitment was further assessed by NeuN co-labeling^72,73^. Approximately 10% and 30% of c-Fos-positive cells in the IC were NeuN-positive in the small- and large-craniotomy conditions, respectively, whereas ∼98% of c-Fos-positive cells in A1 were NeuN-positive in both conditions. Thus, despite the lack of clear electrophysiological evidence for direct cortical activation, c-Fos/NeuN labeling indicates neuronal recruitment within the CNIC and A1, with the marked difference in NeuN-positive fractions suggesting distinct modes of recruitment: FUS may locally engage both neuronal and non-neuronal cells in the IC, whereas cortical activation is more likely relayed synaptically through the auditory afferent pathway. Notably, the ∼3-MPa fundamental pressure reaching the IC was below the range that robustly activated neurons in our 20-MHz in-vitro DRG^25^ and SGN^26^ experiments, although those studies used shorter 1-ms pulses. FUS may therefore also recruit mechanosensitive non-neuronal cells, such as astrocytes, which could indirectly engage neighboring neurons through gliotransmitter signaling^74,75^.

In contrast, c-Fos expression in the DCN, predominantly localized to the granule-cell region, did not differ between ipsilateral and contralateral sides, nor markedly between FUS and control conditions, arguing against substantial recruitment of the cochlear nucleus by IC-targeted FUS. We also observed few to no c-Fos-positive cells in the ventral cochlear nucleus. The cochlear-nucleus granule-cell domain receives prominent non-auditory inputs from the cuneate and spinal trigeminal nuclei, as well as vestibular pathways, integrating somatosensory and proprioceptive information related to head and pinna position^72–74^. Thus, the sparse, comparable DCN labeling across conditions and sides is more consistent with ongoing background multisensory activity than with FUS-induced activation.

Because c-Fos can also be induced by cellular stress and neuronal injury^76^, we considered whether the observed labeling reflected FUS-induced tissue injury. This possibility is unlikely: cleaved caspase-3 immunoreactivity was not increased after FUS, and three-dimensional bioheat simulations predicted a maximum temperature rise of only ∼0.7–0.8°C in the targeted IC, well below levels associated with thermal brain injury^77^. Similar c-Fos and cleaved caspase-3 analyses have been used previously to distinguish ultrasound-driven cellular recruitment from apoptotic damage^78^. Together with the spatial confinement of c-Fos labeling to the FUS target and NeuN co-labeling, these findings support FUS-driven central cellular recruitment rather than injury-associated c-Fos expression.

The increase in the number of c-Fos-positive cells in A1 for the SC and LC conditions suggests that FUS stimulation of the IC can activate the afferent pathways without inducing short-latency, well-synchronised neural responses in the auditory cortex, as indicated by the absence of large-amplitude, reliable LFP’s in the LC condition. The reason for this is unclear. One possibility is that FUS stimulation does not produce the same fast, synchronized neural responses as electrical stimulation, which are required to activate the ascending pathway^62^. FUS stimulation may potentially activate neurons via slower mechanisms, such as the one described above involving calcium signaling amplification with TRPP1/2 and TRPC1 channels. Stimulation of the FUS may also slowly modulate neuronal activity in the CNIC via astrocytes and their TRPA1 channels. This process of neural stimulation has been reported to take several seconds^75^. In conclusion, it is conceivable that FUS stimulation used in this study slowly modulates neural activity instead of triggering fast, synchronized responses.

Moreover, our study also shows convincingly that FUS targeting the IC can stimulate efferent pathways, possibly also via the mechanisms developed above. The MOC system is primarily a cochlear gain modulation system, rather than a sensory activation system that faithfully encodes acoustic information particularly temporal information^79^. Therefore, assuming that the MOC system can be activated by slow activity originating from the IC, FUS stimulation could also activate the descending pathway and impact the cochlea. We demonstrate a slow, cumulative effect of FUS stimulation on sound-evoked cochlear potentials. Specifically, the amplitude of sound-evoked CAP responses decreased, while the amplitude of CM responses increased. These results were observed in both small- and large-craniotomy conditions and were particularly pronounced in the small-craniotomy condition at 8 kHz and at low levels. The effects were still evident an hour after the stimulation sequence. The MOC system likely plays a key role in these results, as gentamicin-mediated blockade of MOC neurotransmission was found to eliminate the changes in CAP and CM amplitudes induced by FUS^68^. These results are reminiscent of the “slow effect” of efferent stimulation on cochlear potentials^65^. The “slow effect” on the cochlea (a decrease in CAP amplitude and an increase in CM amplitude recorded 600 ms after electrical stimulation of the olivo-cochlear bundle) is mediated by the MOC system and builds up over time while the olivo-cochlear bundle is repeatedly electrically stimulated (with electric shocks being administered every 1.5 seconds). Interestingly, the “slow effect” on the cochlea can persist for tens of seconds after the final electric shock. The “fast effect”, on the other hand, occurs immediately after each electric shock and does not change over time^65^. When the MOC system is activated, it releases acetylcholine at the synapses with the outer hair cells (OHCs). This elicits calcium entry into the OHCs through alpha9/10 receptors, increasing the potassium conductance through the associated SK2 channels. Ultimately, this leads to potassium exit, hyperpolarization of the OHCs, and a decrease in their contractile properties. The reduction in CAP response amplitude is directly related to the decrease in OHC contribution to cochlear amplification. Furthermore, the increase in potassium conductance through SK2 channels can account for the increase in CM amplitude^65^. In guinea pigs, it has been shown that the descending IC projection to the superior olivary complex arises primarily from the central nucleus, secondarily from the dorsal cortex and finally from the external cortex^80^. It is therefore entirely plausible that FUS activates the efferent neurons of the dorsal cortex and the central nucleus of the IC (see above the paragraph on spatial dimension of the FUS beam).

Several biophysical mechanisms have been proposed to explain FUS-induced neurostimulation, including thermal effects, cavitation, and mechanical effects. Under our experimental conditions, the thermal simulations predicted a maximum temperature rise of approximately 0.8°C at the focal spot in the IC (Figures 8H - J). Although a thermal contribution cannot be entirely excluded, the small temperature increase predicted during a brief 3-ms stimulus makes substantial thermal effects less likely. Indeed, infrared stimulation of the inferior colliculus has been shown to modulate neuronal activity through temperature-sensitive TRP channels, including TRPV1-4 and/or TRPA1, but these effects were observed during much longer, 30-s infrared stimulation^81^. Cavitation is also unlikely to be the primary mechanism under our experimental conditions, as the mechanical index was approximately 1.2, below the commonly used cavitation threshold of 1.9 for soft tissues^82^. Mechanical effects therefore represent a plausible mechanism underlying the neuronal activation observed in our study. These effects may arise from two distinct processes: the successive bulk compression/dilatation due to the US propagation wave itself, and the acoustic radiation force resulting from momentum transfer from the ultrasound wave to the absorbing or scattering medium. Both types of mechanical perturbation may modulate mechanosensitive signaling pathways and influence neuronal excitability. It has recently been demonstrated that acoustic-pressure-driven mechanical effects, associated with displacements on the nanometer scale, can activate the mechanosensitive RET signalling pathway, demonstrating that FUS can trigger cellular responses through mechanical action independently of acoustic radiation force in tissue^83^. Acoustic radiation force can also produce tissue deformation and mechanical stresses capable of modulating mechanosensitive ion channels and neuronal excitability^84^. Although the relative contributions of these mechanical effects remain to be determined in our experiments, they represent the most likely physical mechanisms underlying FUS-induced activation of the IC.

### Plasticity of the MOC system?

The long-lasting effects of FUS stimulation on the cochlea that were still present one hour after the end of the FUS stimulation sequence, are an interesting and puzzling result. They suggest that FUS stimulation triggered some sort of plasticity mechanisms in the MOC system. Very few studies have reported plasticity of the MOC system, or at least mechanisms compatible with some form of neural plasticity^64,65,67,85^. It has been demonstrated that electrical stimulation of the IC, combined with acoustic stimulation of the contralateral ear (which does not result in any noticeable change in CAP amplitude by itself), leads to a greater reduction in CAP amplitude than IC stimulation alone^67^. Another study showed that olivocochlear efferent neurons were facilitated when both ears were stimulated simultaneously. This binaural facilitation could lower their threshold by as much as 40 dB and increase their discharge rate by as much as 80%. Interestingly, the facilitatory effect could persist even 10 minutes after the end of the binaural stimulation^64^. Recently, the synaptic plasticity of the MOC efferent neurons has also been demonstrated. Repetitive stimulation of the IC can considerably strengthen the synapses of MOC neurons for tens of seconds^85^. Overall, these studies demonstrate that the MOC system has a “reserve” of activation, potentially through plastic mechanisms, that can be harnessed to enhance its role in the auditory system.

I f the MOC system is indeed plastic, a crucial question that must be answered is at what level this plasticity is induced. The precise mechanisms involved in MOC system facilitation, particularly those triggered by FUS stimulation, are largely unknown. The only information we have to help us understand our results is that FUS stimulation activates the cochlea and the outer hair cells (OHCs) via the MOC system within a very short time frame and that cochlear activation is not necessary for MOC system plasticity (as shown by the large craniotomy condition results). Furthermore, our results demonstrate that FUS stimulation provides protection against noise trauma in the SC condition, but not in the LC condition (see below). One possible explanation is that MOC plasticity is enhanced by the simultaneous activation of the IC and of the cochlea. While we do not have statistical evidence for a stronger increase in CM and decrease in CAP in animals with a small craniotomy compared to animals with a large craniotomy, our data exhibit a trend in this direction (e. g., −0. 97 ± 0. 1 5 dB CAP decrease for LC vs. −2. 4 ± 0. 27 dB for SC at 8 kHz). CAP recordings show that FUS-related cochlear nerve activation occurs within 1 ms. The cochlear nucleus, MOC neurons and inferior colliculus should therefore be activated at 2, 3 and 6 ms after FUS stimulation, respectively. Taking into account the stimulation of efferent neurons in the IC and the propagation of this activity, the MOC neurons in the superior olivary complex are expected to be activated within 3 ms of FUS stimulation. Ultimately, hyperpolarization of outer hair cells (OHCs) should therefore occur around 7 ms after FUS stimulation.

From these approximate values, we can infer that FUS-related cochlear activation likely occurs a few milliseconds before OHC modulation by MOC fibers. Could internal changes within the OHCs, related to the activation of the OHCs just before a modulation of the MOC system, fully account for our results? We cannot answer this question with certainty, but this mechanism seems unlikely to us, especially as an explanation for the protective mechanisms against auditory trauma. Indeed, activation of the MOC system via electrical stimulation reduces hearing loss by only 1 0–20 dB, which is far from the 60 dB we report in this study^1^ ^3, 86^. Another mechanism that may be involved and that could have a powerful effect on the cochlea and its protection against noise trauma is an increase in the MOC system at the central level through plasticity-based mechanisms^82^. Indeed, MOC neurons could act as coincidence detectors for ascending and descending inputs arriving at nearly the same time^87^. This may create the conditions for neural plasticity of MOC neurons, thereby amplifying their contribution to hearing, including the protective effect of the MOC system. Further experiments are needed to identify the mechanisms underlying the plasticity of the MOC system that we observe.

### Functional role of the MOC system plasticity

The exact functional role of this plasticity within the MOC system has yet to be determined. Here, animals that were exposed to a high-level tone for one hour showed a threshold shift of 60 dB at frequencies above 4 kHz. On the other hand, the same acoustic trauma in animals subjected to FUS neuromodulation before the noise trauma showed little or no ABR threshold shift. A protective effect of FUS neuromodulation against noise trauma was observed regardless of whether FUS was applied to the CNIC ipsilaterally or contralaterally to the measured ear. This protective effect was abolished in animals treated with gentamicin, suggesting that the FUS-activated MOC system is involved. Finally, other conditions were tested, including intact skull and large craniotomy conditions. None of these additional conditions showed any protective effect on the cochlea.

Exposure to excessive sound levels can cause various types of damage at the level of the hair cells and stereocilia^88,89^, the synapse between hair cells and the cochlear nerve (synaptopathy^90^) and the cochlear nerve (degeneration)^91^. While the underlying cellular mechanisms of NIHL are not fully understood, damage can include the mechanical disruption of stereocilia^92^ and/or the accumulation of toxic molecules (such as glutamate, free radicals and potassium) within hair cells, which can lead to apoptosis^89,91,93,94^. It is important to bear in mind that noise trauma can cause temporary or permanent damage to the cochlea. This means that the auditory system has a natural ability to repair itself following noise trauma^92,93,95–98^. For instance, several hair cell structures (stereocilia, actin cores, tip links and rootlets) have been shown to undergo partial repair following trauma^92,98^. However, it seems that only certain types of damage can be repaired after auditory trauma, despite normal thresholds being restored^89,98^. For instance, moderate noise trauma causing complete reversible threshold elevation can lead to an uncoupling between IHC and cochlear fibers and delayed degeneration of the cochlear nerve^90^. Moreover, if the damage is too severe, repair is no longer possible, regardless of the type of damage. An important point to discuss here is the extent of FUS’s protective effect against acoustic trauma compared to what has been reported in the past. Many studies have reported that various treatments can reduce hearing loss caused by acoustic trauma^91,93,94,99^. The application of pharmacological agents, including corticoids, antioxidants and neurotrophic factors, before or shortly after noise trauma has been shown to play a protective role^91,99–101^. Other studies have shown that non-pharmacological agents can also have a protective role against noise trauma. For instance, presenting continuous audible sounds at moderate levels prior to noise trauma can reduce hearing loss by up to 30 dB^102,103^. Possible explanations for these effects include strengthening of the MOC system, making the cochlea “tougher” and more resistant to NIHL^104^. Furthermore, the protective effect of the efferent system can be enhanced by stimulating the olivocochlear complex or the IC electrically. The protection of hearing following acute acoustic trauma (60 dB) using FUS stimulation appears to be more effective than most treatments that have already been tested^105^, whether they involve “toughening” (∼25 dB)^102,104^, electrical stimulation or enhancement of the MOC system (∼20 dB)^13,106,107^, or drugs such as corticosteroids (∼15 dB)^105^, antioxidants (∼15 dB)^99^, and neurotrophins (little threshold protection)^108^. Our results place FUS stimulation among the most effective and therefore most promising approaches for protecting hearing.

One remaining question is whether the reduction of cochlear gain via MOC activation is the only mechanism protecting the cochlea from noise trauma in our experiment^79^, or whether there are other mechanisms at play. FUS may play a role in lesions that could be reversible if treated quickly enough after the trauma, provided the correct mechanisms are activated. We can only speculate about these mechanisms. It has recently been shown that the MOC system establishes connections with supporting cells and gap junctions^109^. This mechanism could play a role in protecting hearing following acoustic trauma by improving potassium recycling. Similarly, FUS stimulation, which propagates to the cochlea, may interact with the connexins^110,111^ that form gap junctions^112,113^; by facilitating their opening via mechanical deformation, it could potentially improve potassium recycling and thus limit the associated metabolic damage^114^. Beyond this specific mechanism, FUS stimulation may limit the activation and effects of processes associated with noise overexposure, which would ultimately lead to cell death.

Stimulation of the right IC with FUS may potentially recruit bilateral olivocochlear efferent pathways, including also the lateral olivocochlear (LOC) system. Although LOC neurons predominantly project ipsilaterally to the cochlea, anatomical studies have demonstrated a smaller contralateral component, suggesting that IC stimulation could potentially influence the contralateral cochlea through bilateral olivocochlear pathways^115,116^. Activation of the left LOC could, in turn, modulate auditory-nerve activity and may potentially contribute to protection and recovery of the cochlea against acoustic trauma as shown in a recent study^117,118^.

## Conclusion

This study is the first to demonstrate that focused ultrasound (FUS) can activate neurons in the inferior colliculus in anesthetized animals. Using cochlear electrophysiology (compound action potentials and cochlear microphonics), we show that IC-targeted FUS robustly engages the auditory efferent pathway. Spatially restricted c-Fos expression and c-Fos/NeuN co-labeling in the targeted IC provide evidence for local neuronal recruitment, while c-Fos expression in auditory cortex suggests engagement of ascending auditory circuitry. Notably, we show that activating the efferent system with FUS provides cochlear protection against noise trauma to an extent that has never been observed before. Although further studies are required to understand the mechanisms underlying this protection, this study paves the way for the potential non-invasive use of ultrasound in humans for hearing protection from noise trauma.

## Resource availability Lead contact

Further information and requests should be directed to and will be fulfilled by the lead contact, Arnaud Norena.

## Materials availability

This study did not generate new reagents.

## Data and code availability

Data associated with this article will be deposited in Mendeley Data. Original code will be publicly made available on GitHub.

## Acknowledgments

This work was supported by the Agence nationale de la recherche under grant ANR-19-CE19-0015, by the CNRS Mission pour l’interdisciplinarité (MITI) and by Fondation Pour l’Audition. This work has also benefited from a French government grant managed by the Agence Nationale de la Recherche under the France 2030 program, reference ANR-23-IAHU-0003. The funders had no role in study design, data analysis, the decision to publish, or preparation of the manuscript.

## Author contributions

**Conceptualization,** V.P., E.F., O.M., and A.N.**; Methodology,** V.P., E.F., O.M., and A.N.**; Software,** V.P.**; Investigation,** V.P., G.R., E.F., O.M., and A.N.**; Formal analysis,** V.P., G.R., E.F., O.M., and A.N.**; Resources,** E.F., O.M., and A.N.**; Data curation,** V.P. and G.R.**; Visualization,** V.P. and G.R.**; Writing – original draft,** V.P. and A.N.**; Writing – review & editing,** V.P., G.R., E.F., O.M., and A.N.**; Supervision,** O.M., E.F., and A.N.**; Project administration,** V.P., O.M., and A.N.**; Funding acquisition,** O.M. and A.N.

## Supplemental information

Supplemental information includes seven figures and three tables.

