## Supplementary information 1 for "High-Frequency Focused Ultrasound Directly Activates the Auditory Midbrain and Protects Against Noise-Induced Hearing Loss in Guinea Pigs"

**Supplementary Figures**

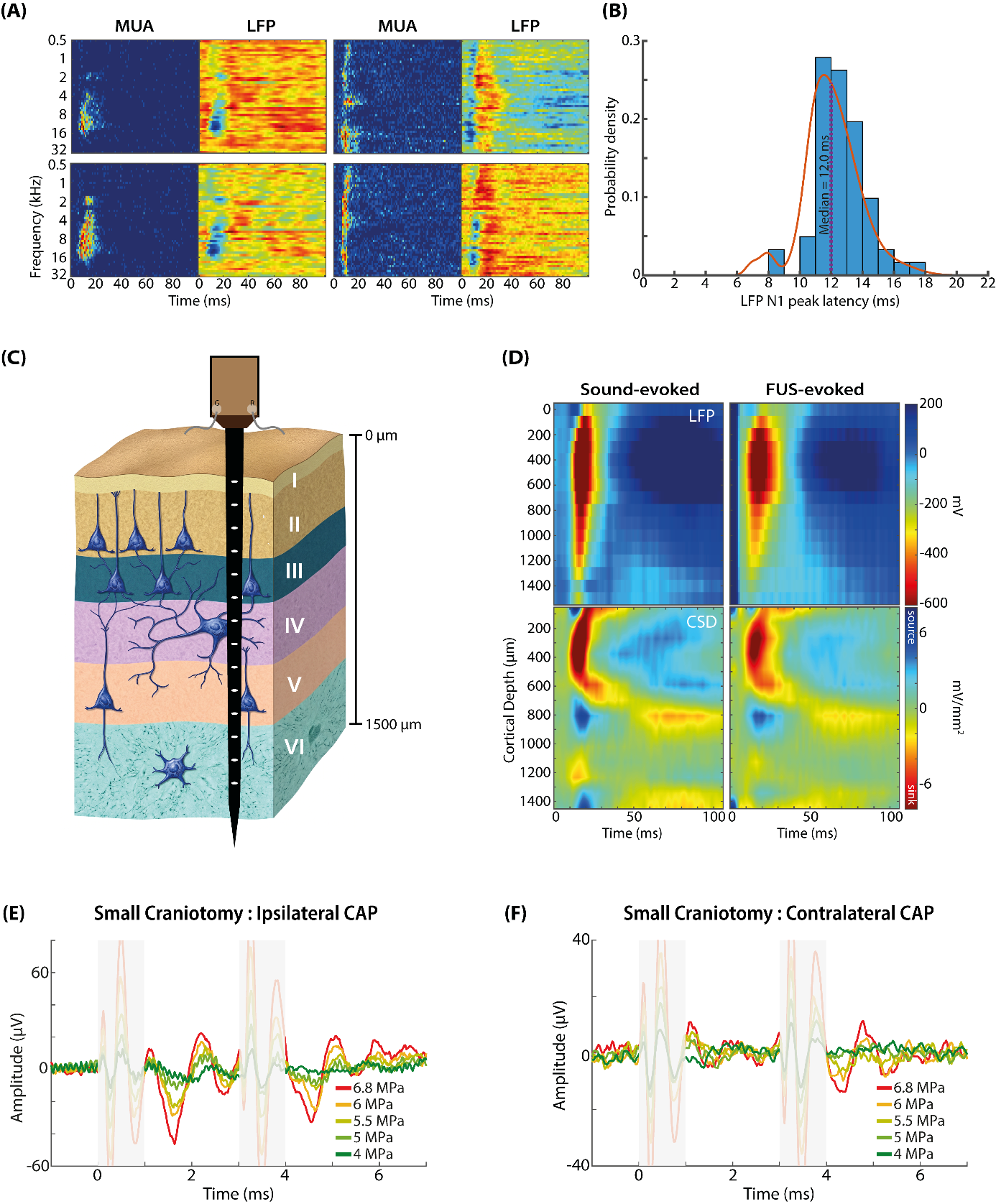

**Figure S1. Electrophysiological characterization of FUS-evoked responses in the central auditory pathway and cochlea.**

(A) Representative multi-unit activity (MUA; left) and local field potential (LFP; right) responses recorded from the central nucleus of the inferior colliculus (CNIC), shown as time-frequency representations of FUS-evoked activity. (B) Probability distribution of LFP N1 peak latencies measured from CNIC recordings (n = 10). The dashed vertical line indicates the median N1 latency. (C) Schematic illustration of the multichannel recording configuration in the auditory cortex, with the linear electrode spanning the cortical layers. (D) Representative laminar LFP (top) and corresponding current source density (CSD; bottom) profiles in the auditory cortex for sound-evoked (left) and FUS-evoked (right) responses. The sound stimulus was presented at the best frequency (BF) of the recorded animal. (E) Representative ipsilateral compound action potential (CAP) responses evoked by FUS delivered through a small craniotomy, shown across increasing ultrasound pressures. (F) Representative contralateral CAP responses recorded under the same stimulation conditions and across the same range of ultrasound pressures.

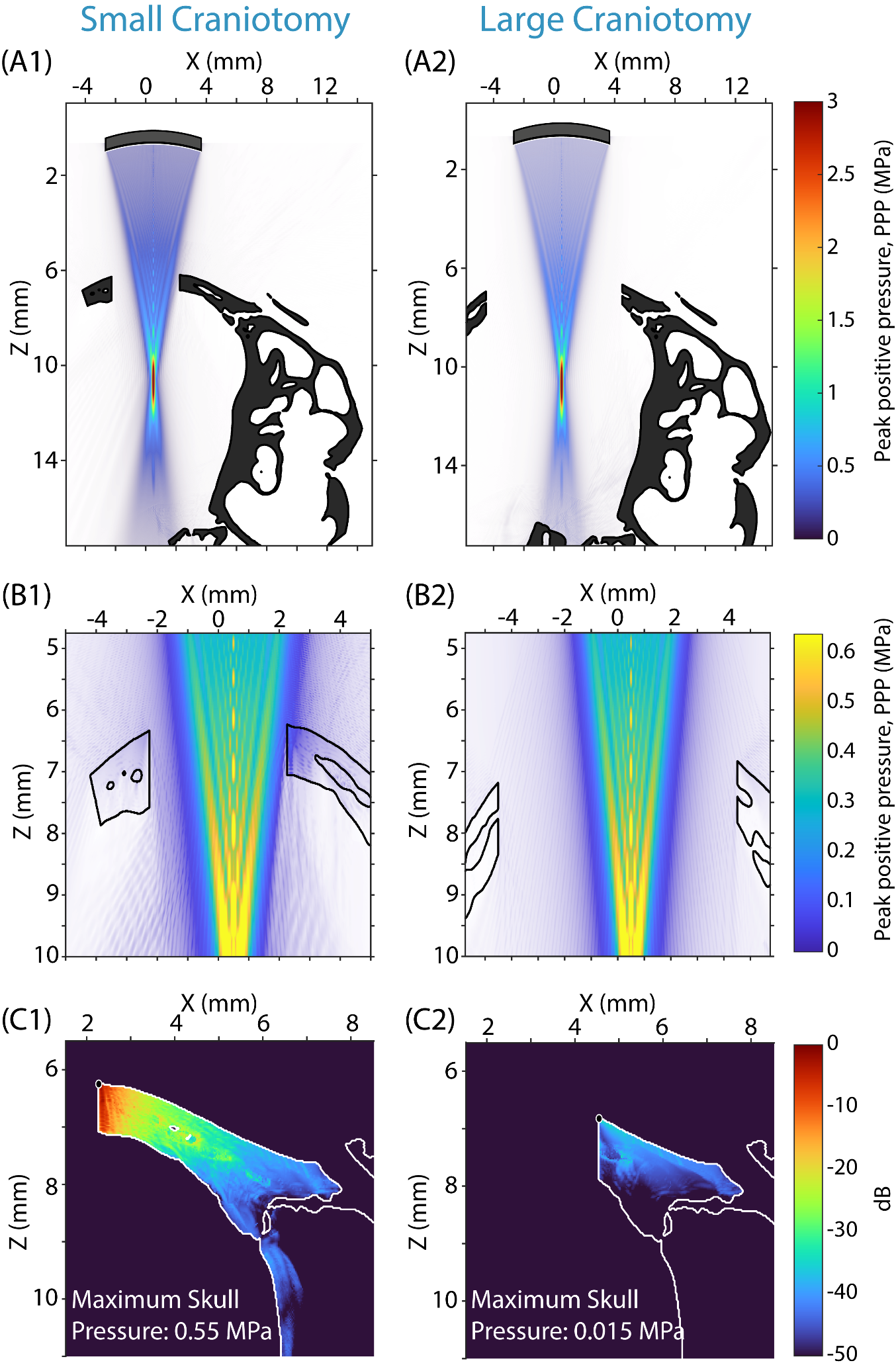

**Figure S2. Acoustic interaction with the skull during FUS stimulation of right IC through small craniotomy (A1, B1, C1) and large craniotomy (A2, B2, C2).**

Acoustic propagation was modeled using k-Wave in a microCT-derived guinea pig skull geometry. (A) Peak-positive pressure (PPP) distribution in the longitudinal view X-Z overlaid on the microCT-derived skull geometry. (B) Enlarged view of the craniotomy region, showing the pressure field within the gel and brain regions as ultrasound propagates through the opening and interacts with the surrounding skull margins. (C) Skull-associated pressure distribution, expressed in decibels relative to the maximum pressure within the skull.

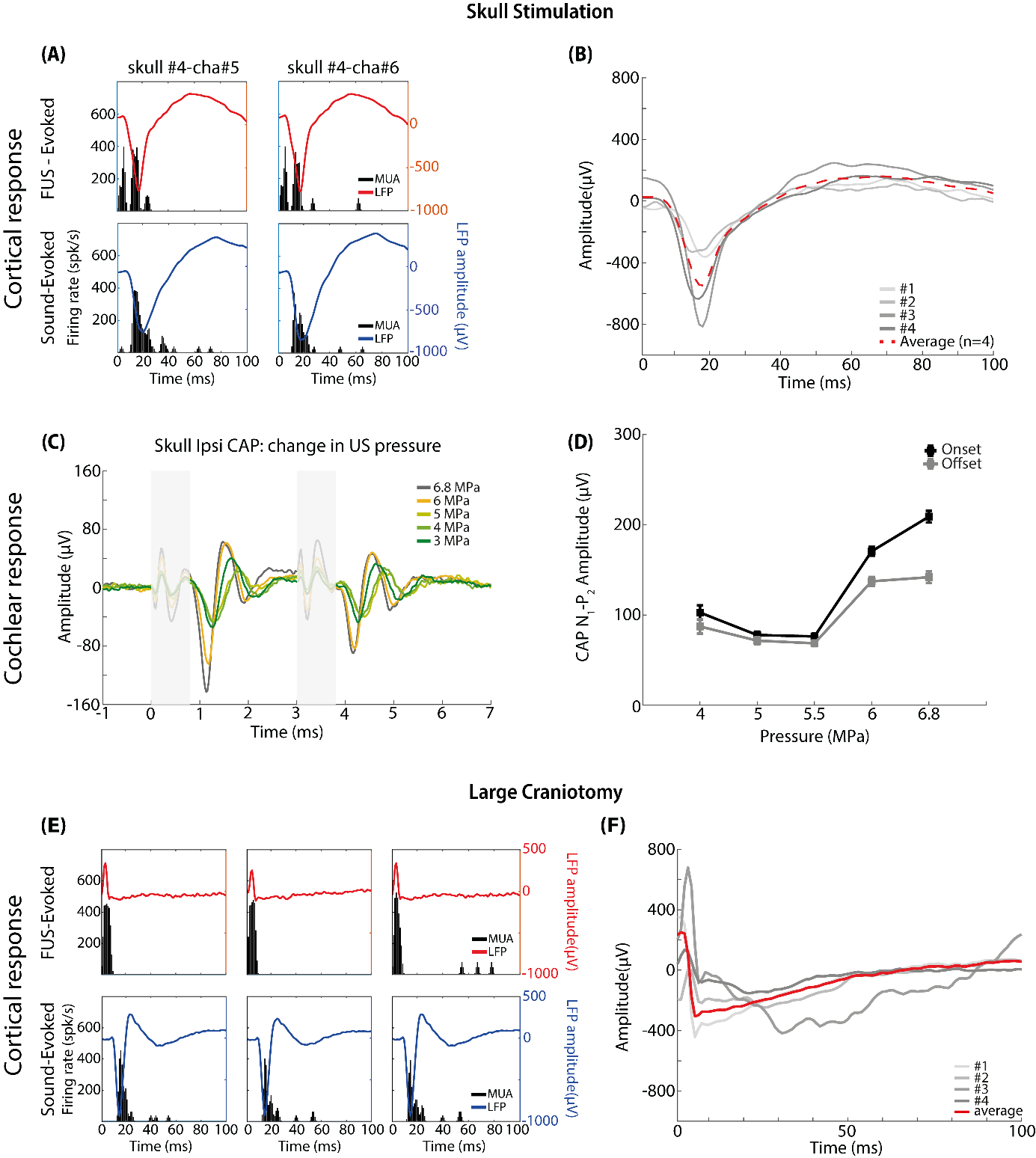

**Figure S3. Cochlear and cortical responses evoked by FUS applied over the intact skull and through a large craniotomy**

(A) Representative layer 4 auditory cortical responses from two recording sites during FUS stimulation applied over the intact skull above the right IC (20 MHz, 6.8 MPa, 3 ms sonication duration, 90 s interstimulus interval [ISI]) and acoustic stimulation at the best frequency (BF) of each recording site (70 dB SPL). Poststimulus time histograms (PSTHs; 1 ms bins) of spiking activity and corresponding LFP responses are shown. LFPs were averaged across 40 FUS trials and 10 sound trials for each recording site. (B) Animal-averaged suprathreshold FUS-evoked layer 4 LFP responses following stimulation over the intact skull. Individual animal averages are shown in gray and the grand average across animals in red. (C) Representative ipsilateral compound action potential (CAP) responses evoked by FUS applied over the intact skull above the right IC, shown across increasing ultrasound pressures. (D) FUS-evoked CAP onset and offset amplitudes as a function of ultrasound pressure during intact-skull stimulation. (E) Representative cortical responses evoked by FUS delivered through a large craniotomy over the IC and by acoustic stimulation. Multi-unit firing rate and simultaneously recorded LFP responses are shown. (F) Individual animal-averaged FUS-evoked cortical LFP responses following stimulation through the large craniotomy (gray) and the corresponding grand-average response across animals (red).

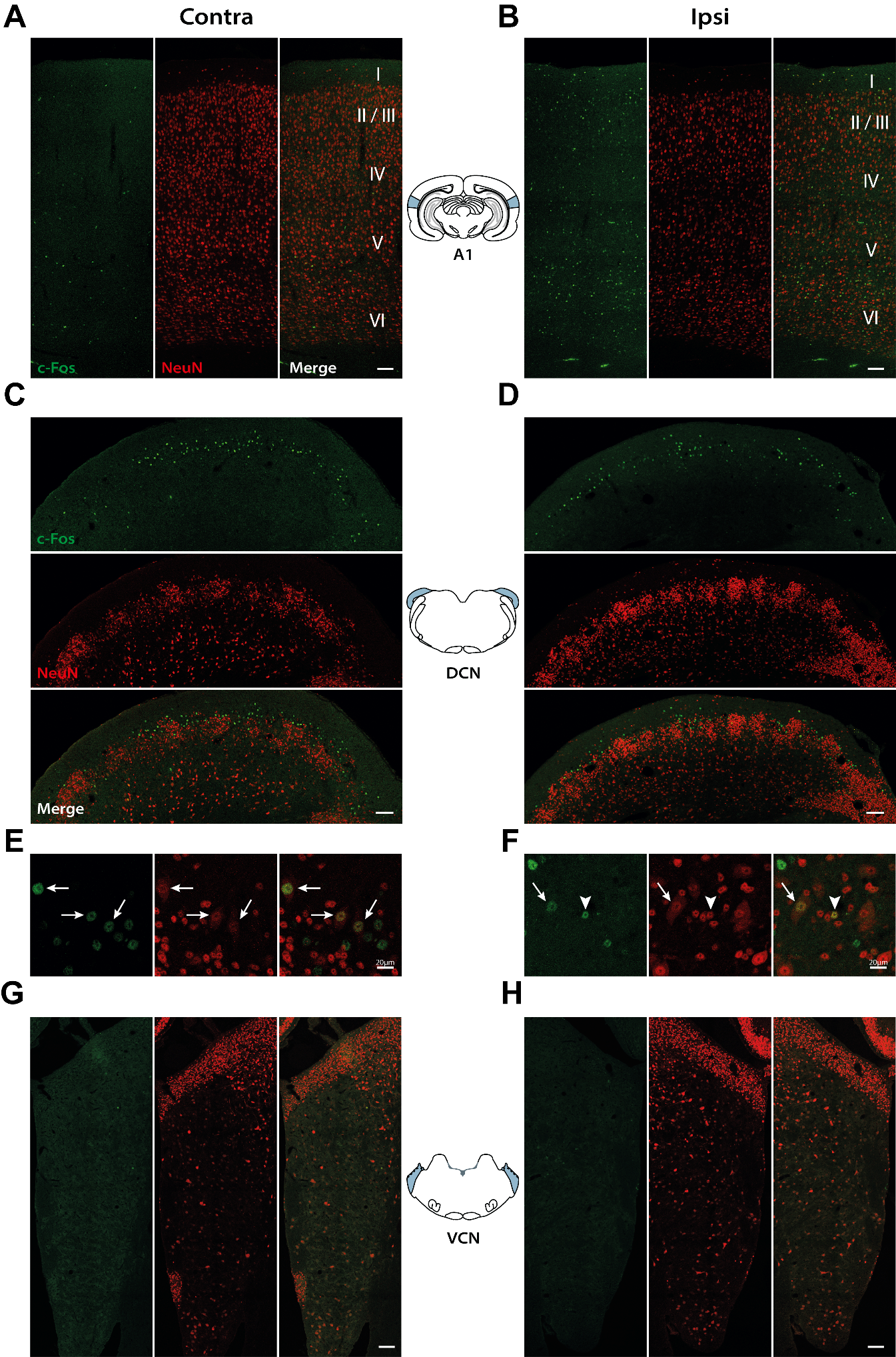

**Figure S4: c-Fos activation in the auditory cortex, DCN, and VCN following IC-targeted FUS.**

(A, B) Representative c-Fos, NeuN, and merged images across cortical layers I–VI of the primary auditory cortex (A1) in the contralateral (A) and ipsilateral (B) hemispheres. (C, D) Representative c-Fos, NeuN, and merged images of the dorsal cochlear nucleus (DCN) in the contralateral (C) and ipsilateral (D) hemispheres. (E, F) Higher-magnification images of the DCN region illustrating that c-Fos/NeuN double-positive cells were predominantly large neurons consistent with fusiform cells (white arrows); occasional c-Fos-positive granule cells were also observed (white arrowhead in F). (G, H) Representative c-Fos, NeuN, and merged images of the ventral cochlear nucleus (VCN) in the contralateral (G) and ipsilateral (H) hemispheres. Scale bars = 100 µm, except in (E) and (F), where scale bars are 20 µm as indicated.

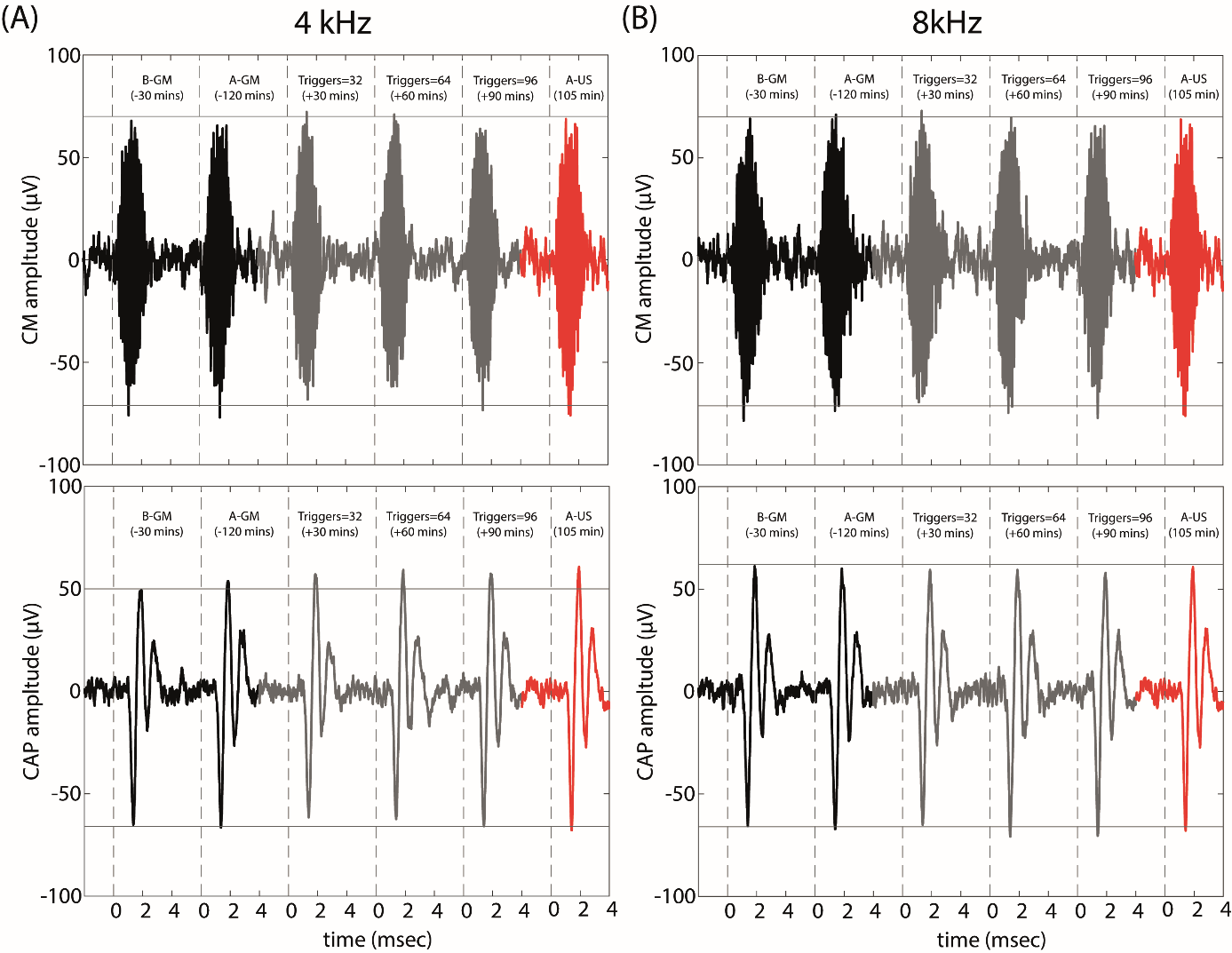

**Figure S5. CAP and CM responses before and after gentamicin treatment during FUS stimulation.**

Representative cochlear microphonic (CM) and compound action potential (CAP) waveforms recorded at 4 kHz (A) and 8 kHz (B) before gentamicin (B-GM), after gentamicin (A-GM), during FUS stimulation at 30, 60, and 90 min, and after FUS (105 min). CM traces are shown above and CAP traces below.

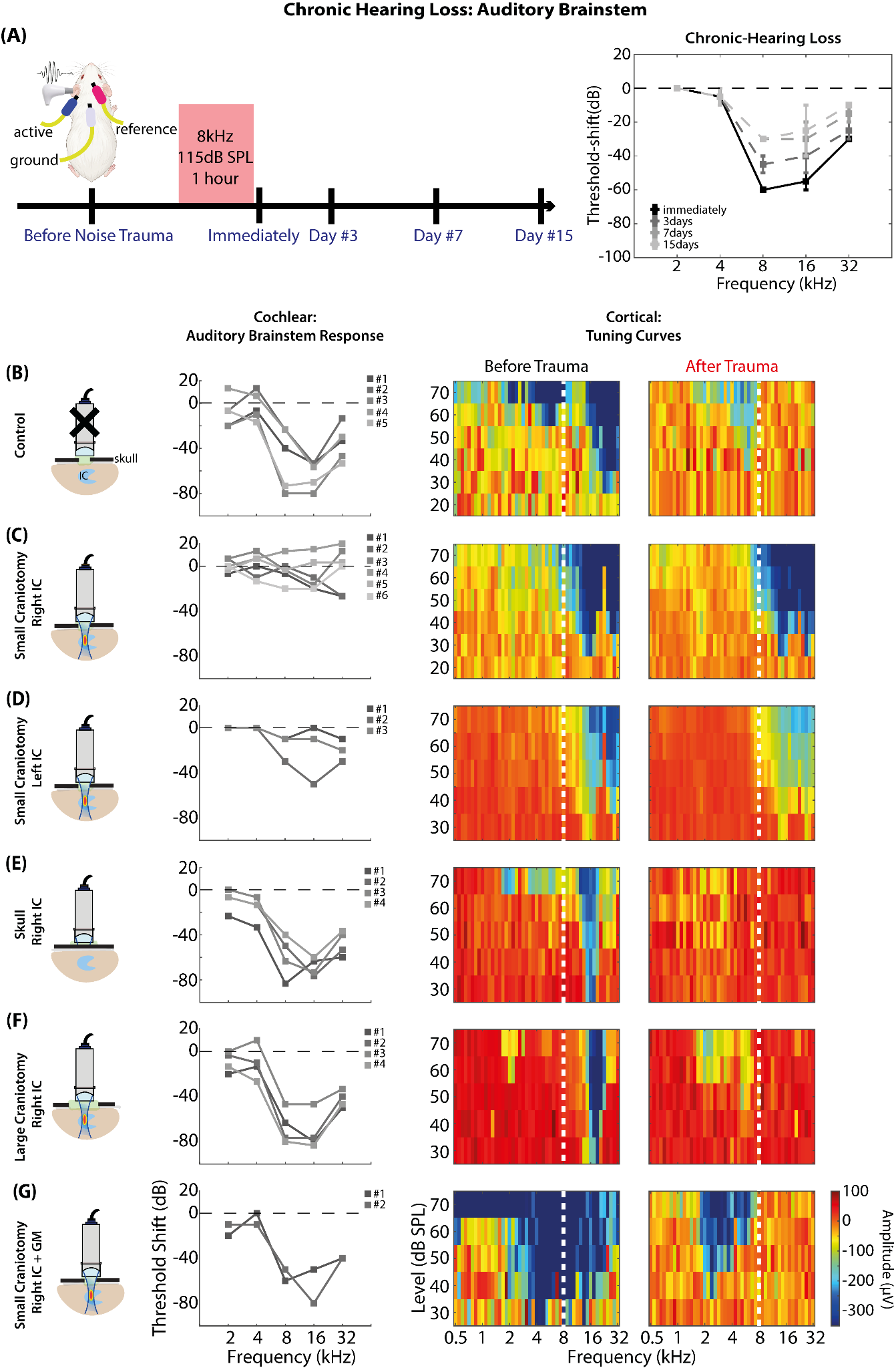
 **Figure S6. Chronic hearing loss, individual ABR threshold shifts, and representative cortical tuning curves across experimental conditions, related to Figure 6 and 7.**

(A) Experimental paradigm for assessment of chronic hearing loss. ABR thresholds were measured before noise exposure and immediately, 3, 7, and 15 days after noise exposure. Right, mean ABR threshold shifts across frequencies at each post-noise time point relative to the pre-noise baseline (n = 3 animals). (B–G) Individual ABR threshold shifts and representative cortical LFP tuning curves for the Control (B), small-craniotomy right-IC FUS (C), small-craniotomy left-IC FUS (D), intact-skull right-IC FUS (E), large-craniotomy right-IC FUS (F), and small-craniotomy right-IC FUS following gentamicin treatment (G) conditions. For ABR threshold shifts, each line represents an individual animal. Right, representative Layer IV LFP tuning curves from a single recording site are shown before and after noise exposure for each experimental condition. LFP responses are plotted as a function of stimulus frequency and sound level, with response amplitude indicated by the color scale. The white dashed line indicates the 8-kHz noise-exposure frequency. SC, small craniotomy; LC, large craniotomy; IC, inferior colliculus; FUS, focused ultrasound; GM, gentamicin; ABR, auditory brainstem response; LFP, local field potential.

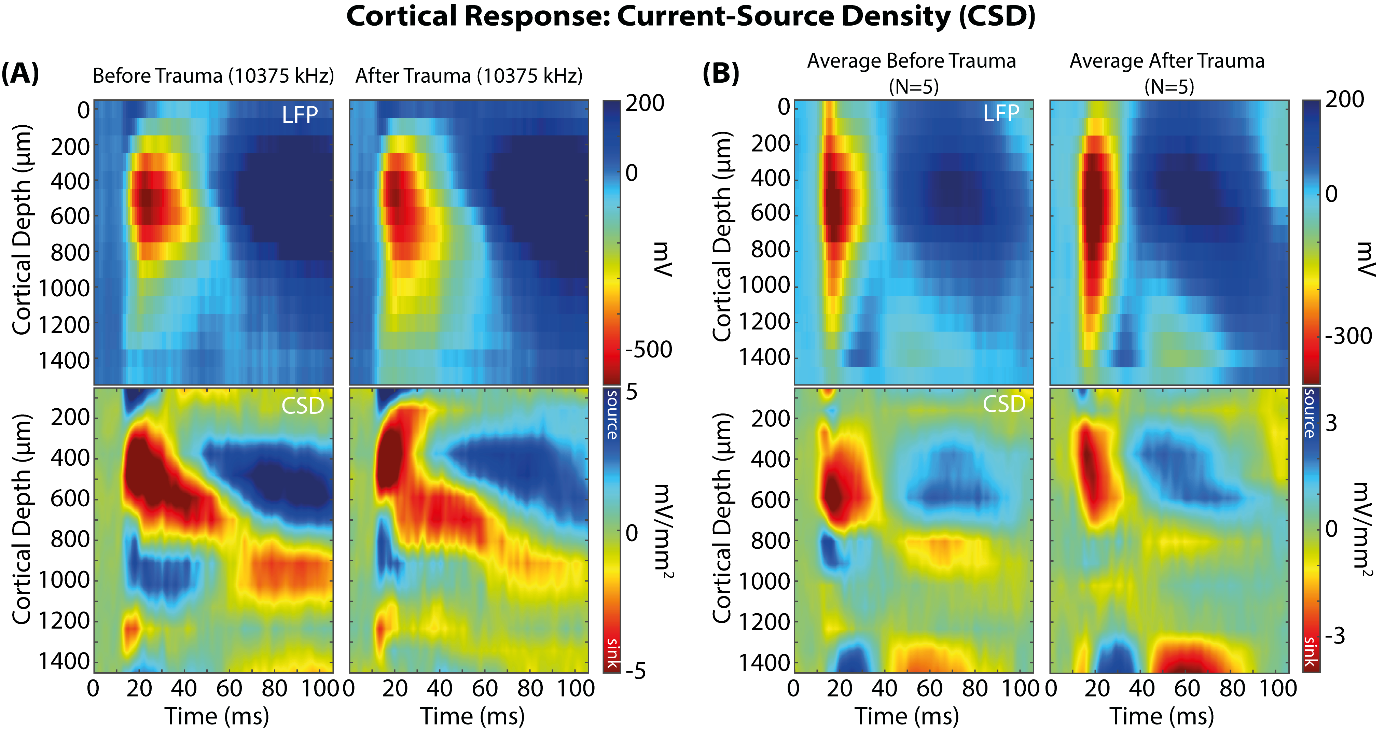

**Figure S7**: **Cortical current source density responses following noise trauma in the small-craniotomy condition**

(A) Representative laminar Local Field Potential (LFP) and corresponding current source density (CSD) profiles of sound-evoked auditory cortical activity before and after NT, measured at the animal’s best frequency (BF). (B) Average laminar LFP and corresponding CSD profiles before and after noise trauma (NT) across five animals.
