## Supplementary information 2 for "High-Frequency Focused Ultrasound Directly Activates the Auditory Midbrain and Protects Against Noise-Induced Hearing Loss in Guinea Pigs"

**Supplementary Tables**

**Table 1**

| Experimental  Condition | N | Animals included in each sub-experiment (N) | Cortical response (%) |
| --- | --- | --- | --- |
| Small Craniotomy: right-FUS | 22 | *Ipsilateral CAP*: 2  *Contralateral CAP*: 6  *Efferent – CAP & CM* : 4  *c-Fos*: 2  *Protection study*: 6  *Gentamicin:* 2 | 79% |
| Small Craniotomy: left-FUS | 3 | *Protection study*: 3 | 55% |
| Large Craniotomy: right-FUS | 9 | *Efferent – CAP & CM efferent*: 3  *c-Fos*: 2  *Protection study*: 4 | 0% |
| Skull: right-FUS | 7 | *US-E Ipsilateral CAP*: 2  *US-E contralateral CAP*: 1  *Protective effect*: 4 | 92% |
| Control: No FUS | 7 | — | — |
| Chronic Hearing loss: No FUS | 3 | — | — |

**T1. Summary of experimental groups**

Animals are grouped according to experimental condition, with the total number of animals (N) indicated for each group. The third column summarizes the number of animals included in each sub-experiment. The final column shows the percentage of Focused Ultrasound (FUS) triggers that elicited a detectable cortical response within each experimental condition. SC, small craniotomy; LC, large craniotomy; FUS, focused ultrasound; CAP, compound action potential; CM, cochlear microphonic.

**Table 2**

| Cochlear Microphonics (CM) | | | | |
| --- | --- | --- | --- | --- |
| Model | **AICc** | **Log-likelihood** | **ΔAICc** | **Akaike weight** |
| Amplitude~Time+Group+Frequency+Intensity | 1624.6 | -784.437 | 0.00 | 0.708 |
| Amplitude~Time*Group+Frequency+Intensity | 1626.7 | -783.237 | 2.31 | 0.223 |
| Amplitude~Time*Group+Time*Frequency+Intensity | 1628.5 | -783.215 | 4.67 | 0.069 |
| Amplitude~Time+Group | 1697.1 | -802.790 | 29.88 | 0.000 |
| Amplitude~Time*Group*Intensity+Frequency | 1708.6 | -813.320 | 33.45 | 0.000 |
| Compound Action Potential (CAP) | | | | |
| Model | **AICc** | **Log-likelihood** | **ΔAICc** | **Akaike weight** |
| Amplitude~Time+Group+Frequency+Intensity | 1624.6 | -801.450 | 0.00 | 0.673 |
| Amplitude~Time*Group+Frequency+Intensity | 1626.7 | -801.343 | 2.14 | 0.231 |
| Amplitude~Time*Group+Time*Frequency+Intensity | 1628.5 | -801.024 | 3.90 | 0.096 |
| Amplitude~Time+Group | 1697.1 | -840.025 | 72.54 | 0.000 |
| Amplitude~Time*Group*Intensity+Frequency | 1708.6 | -851.760 | 77.17 | 0.000 |
| Auditory Brainstem Responses: threshold-shift | | | | |
| Model | **AICc** | **Log-likelihood** | **ΔAICc** | **Akaike weight** |
| Thresholds~Frequency*Time+Time*Group+  Frequency*Group | 1795.1 | -854.267 | 0.00 | 0.936 |
| Thresholds~Time*Group*Frequency | 1800.4 | -831.720 | 5.35 | 0.064 |
| Thresholds~Frequency*Time+Frequency*Group | 1867.0 | -895.837 | 71.88 | 0.000 |
| Thresholds~Frequency+Time+Group | 1931.6 | -953.029 | 136.47 | 0.000 |
| Thresholds~Time*Group | 2044.4 | -1009.434 | 249.28 | 0.000 |
| Auditory Brainstem Responses: Wave III amplitudes | | | | |
| Model | **AICc** | **Log-likelihood** | **ΔAICc** | **Akaike weight** |
| Amplitude~Frequency*Time+Frequency~Group+  Time*Group+Intensity+ (Group\|AnimalID) | 460.9 | -199.955 | 0.00 | 0.713 |
| Amplitude~Frequency*Time+Time*Group+  Intensity+ (Group\|AnimalID) | 463.1 | -205.372 | 2.18 | 0.240 |
| Amplitude~Frequency*Time*Group+Intensity+  (Group\|AnimalID) | 466.9 | -198.39 | 5.98 | 0.036 |
| Amplitude~Frequency+Time+Group+Intensity+  (Group\|AnimalID) | 470.2 | -214.35 | 9.26 | 0.007 |
| Amplitude~Time*Group+Intensity+ (Group\|AnimalID) | 471.1 | -212.116 | 10.13 | 0.004 |

**T2. Mixed-effects model comparison and model selection for cochlear and auditory brainstem response measurements.**

Candidate mixed-effects models were compared for four outcome measures: cochlear microphonic (CM) amplitude, compound action potential (CAP) amplitude, auditory brainstem response (ABR) threshold shift, and ABR Wave III amplitude. For each dataset, alternative models containing different combinations of the experimental factors and their interactions were fitted and compared using the corrected Akaike information criterion (AICc). Model performance is reported as the AICc, log-likelihood, ΔAICc relative to the best-supported model, and Akaike weight. The model with the lowest AICc (ΔAICc = 0) was considered the best-supported model, while Akaike weights indicate the relative support for each candidate model within the model set. Models were ranked independently for each outcome measure.

**Table 3**

| **Parameter** | **Water calibration** | **Full CT propagation** | **3D thermal model** |
| --- | --- | --- | --- |
| *COMPUTATIONAL IMPLEMENTATION* | | | |
| **Software / solver** | MATLAB; k-Wave v1.4.0; kspaceFirstOrder3D / OMP | MATLAB preprocessing; standalone kspaceFirstOrder -OMP | MATLAB custom 3D bioheat solver |
| **Computational platform** | CPU/OpenMP | Linux HPC; 64 OpenMP threads; high-memory node | High-memory CPU/MATLAB |
| *TRANSDUCER AND SOURCE* | | | |
| **Transducer model** | Olympus V317-SM geometry | Same | Acoustic field imported |
| **Center frequency** | 20 MHz | 20 MHz | 20-MHz acoustic field |
| **Aperture diameter** | 6.32 mm | 6.32 mm | — |
| **Radius of curvature** | 12.7 mm | 12.7 mm | — |
| **Source representation** | kWaveArray off-grid spherical bowl | kWaveArray off-grid spherical bowl | — |
| **BLI tolerance / upsampling** | 0.05 / 10 | 0.05 / 10 | — |
| **Source pressure amplitude** | 0.160 MPa | 0.160 MPa | — |
| **Waveform** | Rectangular sinusoid; no ramp | Rectangular sinusoid; no ramp | 3-ms heating interval |
| **Burst length** | 20 cycles | 50 cycles | — |
| *SPATIAL / NUMERICAL DISCRETIZATION* | | | |
| **Acoustic grid spacing** | 12.5 µm isotropic | 12.5 µm isotropic | 25 µm isotropic |
| **Grid size** | 640 × 640 × 1200 voxels | 1701 × 960 × 1944 voxels | Derived from CT acoustic volume |
| **Perfectly matched layer** | 0.50 mm | 0.50 mm (40 voxels) | — |
| **CFL** | 0.149 | ≤0.15 | — |
| *ACOUSTIC MATERIAL PROPERTIES* | | | |
| **Coupling / water: c** | 1490 m/s | 1490 m/s | — |
| **Coupling / water: ρ** | 1000 kg/m³ | 1000 kg/m³ | — |
| **Brain: c** | — | 1500 m/s | — |
| **Brain: ρ** | — | 1000 kg/m³ | 1000 kg/m³ |
| **Skull: c** | — | 2400 m/s | 2400 m/s |
| **Skull: ρ** | — | 1850 kg/m³ | 1850 kg/m³ |
| **Nonlinearity, B/A** | 4.96 | 4.96 for all fluid-equivalent materials, 6.7 for tissue/Brain | — |
| **Attenuation exponent** | 1.18 (CT-matched representation) | 1.18 globally | 1.18 for brain/skull |
| **Water attenuation** | 0.022161451 dB/(cm·MHz^1.18) | 20-MHz-matched equivalent of native water attenuation | — |
| **Brain attenuation** | — | 0.21 dB/(cm·MHz^1.18) | 0.024 Np/(cm·MHz^1.18) |
| **Skull attenuation** | — | 2.70 dB/(cm·MHz^1.18) | 31 Np/(m·MHz^1.18) |
| *RECORDED / DERIVED ACOUSTIC METRICS* | | | |
| **Recorded pressure outputs** | p_raw, p_max, p_min in focal patch | Whole-volume p_max_all and p_min_all; p_raw at focal/cochlear/route sensors | Acoustic field used as thermal source |
| **Harmonic analysis** | A1 (20 MHz), A2 (40 MHz) | A1/A2 at selected raw-pressure sensors | — |
| **Steady-state analysis window** | 10 cycles (cycles 6–15); no FFT taper | Steady raw-pressure window at selected sensors | — |
| **3D THERMAL MODEL** | | | |
| **Governing equation** | — | — | $\rho C_{p}\frac{\partial T}{\partial t}=\nabla\cdot\left( k\nabla T \right)+Q_{\mathrm{US}}+\rho_{b}C_{b}\omega_{b}\left( T_{b}-T \right)$ |
| **Acoustic heat source** | — | — | $Q_{\mathrm{US}}=\alpha p^{2}/Z$ |
| **Brain heat capacity, Cp** | — | — | 3600 J·kg⁻¹·K⁻¹ |
| **Brain thermal conductivity, k** | — | — | 0.528 W·m⁻¹·K⁻¹ |
| **Skull heat capacity, Cp** | — | — | 1300 J·kg⁻¹·K⁻¹ |
| **Skull thermal conductivity, k** | — | — | 0.40 W·m⁻¹·K⁻¹ |
| **Brain perfusion** | — | — | 0.008 s⁻¹ |
| **Heating duration** | — | — | 3 ms |
| **Post-FUS cooling** | — | — | 200 ms |
| **Thermal boundary condition** | — | — | Zero normal heat flux at outer boundaries |

**T3: Numerical and material parameters for 3D acoustic and thermal modeling**

Summary of parameters used for water calibration, nonlinear micro-CT acoustic propagation, and 3D thermal modeling. Acoustic simulations were performed with k-Wave using a 20-MHz kWaveArray transducer model, and thermal calculations used the simulated pressure field and CT-derived material segmentation. Values correspond to the final simulations used for the manuscript figures.
